# Evidence for holocentric centromeres in the early branching apicomplexan parasite *Cryptosporidium parvum*

**DOI:** 10.1101/2025.09.30.679541

**Authors:** Abigail Kimball, Wanyi Huang, Rui Xu, Melanie Key, Lisa Funkhouser-Jones, L. David Sibley

## Abstract

During its asexual cycle, *Cryptosporidium parvum* (*C. parvum*) amplifies through three rounds of nuclear division before undergoing cytokinesis to form infectious merozoites. However, chromosome organization and segregation during nuclear division remain unstudied. Here, we visualized H3-histones, including the centromeric histone H3 (CENH3), telomeres, and centrosomes during mitosis. Nuclear division was accompanied by centrosome duplication and elongated microtubules that spanned the dividing nuclei. Surprisingly, *C. parvum* centromeres detected by CENH3 showed diffuse staining throughout nuclear division, that overlapped with telomeres in the apical nucleus. Analysis of centromeres based on CENH3 capture and DNA sequencing revealed numerous distinct binding sites scattered across all eight chromosomes, including both coding regions and intergenic regions, that were typified by GA rich repeat sequences. The unique holocentric chromosome structure of *C. parvum* is unlike the single regional centromeres found in related apicomplexans, suggesting it arose independently from other known examples in plants, insects and worms.

## INTRODUCTION

*Cryptosporidium parvum (C. parvum)*, an obligate intracellular parasite, remains a serious global health and agricultural threat [1–4]. Transmission occurs through the ingestion of infectious oocysts and causes severe diarrheal illness in infants, young animals, and immunocompromised adults [5–7]. Once ingested, sporozoites exit the oocyst, invade host cells, and undergo three rounds of nuclear division within a single shared cytoplasm (the process of merogony), followed by cytokinesis to produce eight identical merozoites. Newly formed merozoites are released into the lumen, where they invade new host cells, and repeat the invasion-egress cycle three times before transitioning to the sexual phase of the life cycle [8, 9]. Despite its significance as a global health problem, and the novelty of apicomplexan nuclear division processes, *C. parvum* mitosis remains largely uncharacterized, with our current understanding limited to basic microscopy studies [8, 10–15].

To divide rapidly, apicomplexans appear to rely on several features that are thought to streamline nuclear division. Comparative cell biology suggests that maintaining chromosomes in a minimally condensed state may reduce the need for extensive chromatin remodeling during division, although the extent to which this directly accelerates mitosis remains to be fully resolved [16, 17]. This adaptation necessitates strict nuclear organization during mitosis, where centromeres are typically maintained on the apical end of the nucleus attached to the centrosomes, and telomeres persistently cluster at the basal end of the nuclear periphery [18–20]. Additionally, apicomplexans maintain their nuclear envelope throughout mitosis, minimizing the energy and time required for breakdown and reformation [17, 21]. A consequence of this closed mitosis is restricted access of spindle fibers to chromosomes, necessitating alternative segregation mechanisms based on intranuclear spindles. Rather than centrosomes orchestrating microtubule organization from the cytoplasm, apicomplexan centrosomes (spindle pole bodies, microtubule-organizing centers) are embedded within the nuclear envelope and serve as spindle assembly hubs that elongate the microtubules so they can attach to the centromeres and ultimately coordinate chromosome movement during division [22–29]. It has been postulated that nuclear pores are essential for nucleocytoplasmic transport through the intact envelope and directly support the process of spindle assembly [30, 31]. Electron microscopy studies suggest that *C. parvum* undergoes schizogony, with multiple rounds of nuclear replication preceding a cytoplasmic division, followed by the egress of daughter parasites from the host cell [13].

During mitosis and meiosis, centromeres function to orchestrate chromosome segregation by serving as sites for microtubule anchoring to assure faithful separation at metaphase [32]. Centromeres have variable architecture in different organisms consisting of point centromeres (small 250 bp single site per chromosome) in yeast [33], regional centromeres (gene poor islands of heterochromatin ranging from several to hundreds of kb) in *T. gondii* and *P. falciparum* [26, 27, 34], and holocentric chromosomes (multiple centromere sites along the chromosome) found in multiple species of plants, insects and worms [35–37]. Although centromere function is essential and highly conserved across eukaryotes, centromeric DNA sequences and size are subject to rapid evolution [38–40]. Variation in centromere architecture affects nuclear organization, karyotype changes, and ultimately drives speciation [41, 42].

The nuclear structures, chromosome features, and mitotic mechanisms, including the role of centromeres, have not been examined previously in *C. parvum*. Here, we delineate the fundamental biology of *C. parvum* mitosis, revealing significant deviations from previously assumed mechanisms. Using high-resolution imaging we examined nuclear division during the first round of merogony, leveraging multiple transgenic parasite strains and mitosis-specific antibodies. Additionally, we used Cleavage Under Targets and Nuclease Release (CUT&RUN) [43] to identify the location of the centromeres based on CENH3 and patterns of centromere-associated histone modifications in *C. parvum* and the related parasite *Toxoplasma gondii* (*T. gondii*). In contrast to the monocentric centromeric structure of *T. gondii*, we found that *C. parvum* has a dispersed centromeric chromosome structure with CENH3 binding sites scattered across multiple small regions on all 8 chromosomes, resulting in diffuse CENH3 centromere staining throughout mitosis. Overall, these results suggest that *C. parvum* employs a divergent mitotic mechanism compared to other well-studied apicomplexans and has convergently evolved holocentric architecture.

## RESULTS

### Characterization of centromeric histone H3 (CENH3) and histone H3 candidates in C. parvum

The genome annotation of *C. parvum* contains two genes encoding Histone H3 *cgd3_2540* and *cgd4_3220* (referred to here as H3.1 and H3.2, respectively) and a third gene *cgd4_2030* annotated as a centromere-specific Histone H3 variant (CENH3). To validate these annotations, we performed multiple BLASTP analyses using the protein products of each of these three *C. parvum* genes to identify top hits in the following groups: 1) Apicomplexa (excluding all *Cryptosporidium* species), 2) all eukaryotic organisms (excluding all Apicomplexa), 3) *Homo sapiens*, and 4) *Saccharomyces cerevisiae.* Collectively, these analyses provide strong support for cgd4_2030 as the CENH3 ortholog in *C. parvum* (**Table 1**). To examine the relationship between the protein sequences for the *C. parvum* genes of interest and the top hits of BLASTP analyses, a phylogenetic tree was generated (**Fig 1A**). We found that *cgd3_2540* (H3.1) and *cgd4_3220* (H3.2) closely cluster with histone H3 sequences from other organisms, whereas *cgd4_2030* (CENH3) clustered with other centromeric histone H3 proteins. We then directly compared the amino acid sequences of H3.1 and H3.2 with a global sequence alignment and found that the two sequences were highly similar (99.26%), but less similar compared to the CENH3 (65.10% and 64.43%, respectively) (**Table 2 and Fig 1B**).

**Figure 1:**
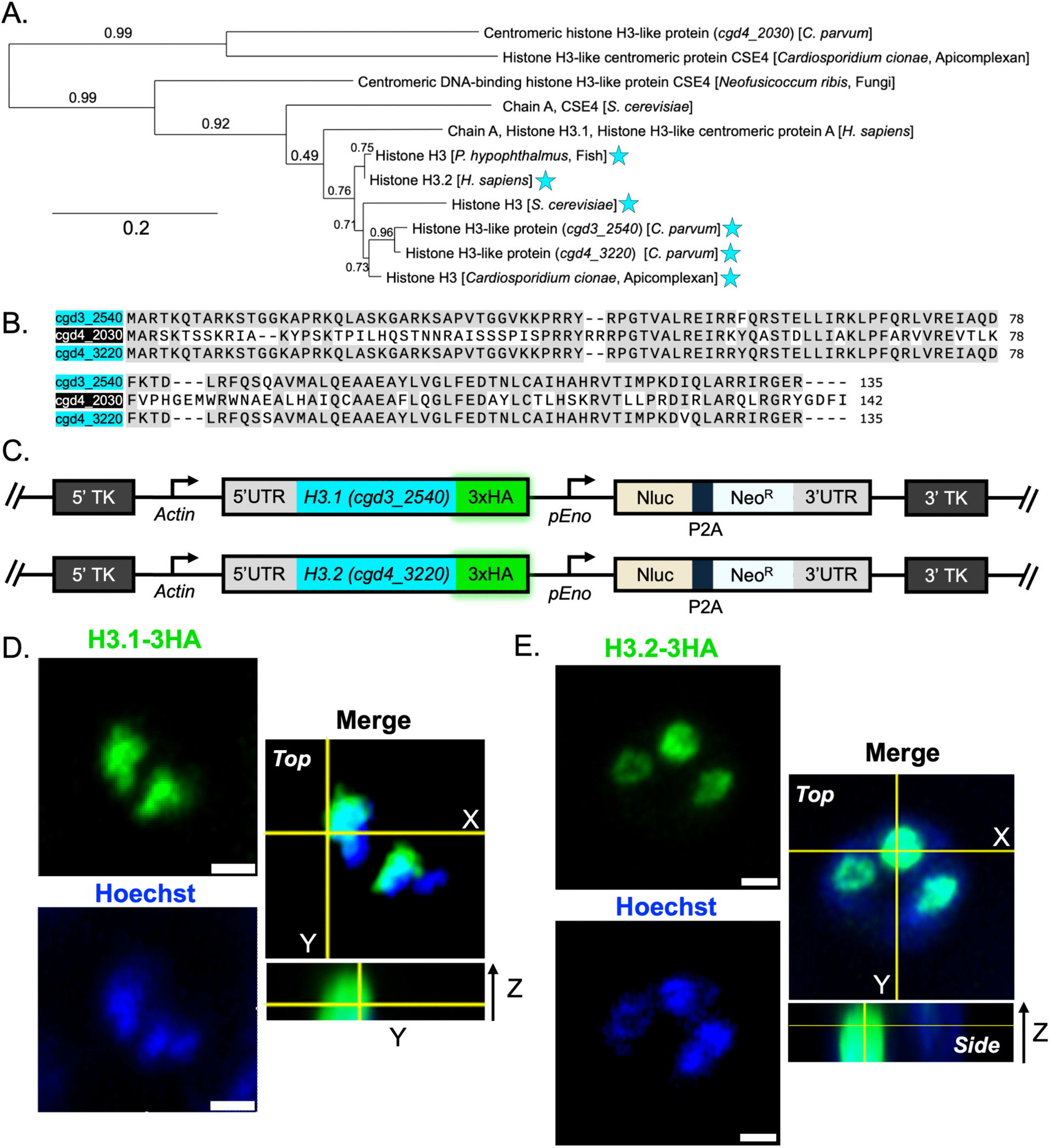
Characterization of centromeric histone H3 (CENH3) and histone H3 candidates in *C. parvum* and construction of Histone H3-3HA tagged strains. (A) Phylogenetic relationship of *C. parvum* H3 and centromeric H3 (CENH3) histones and their corresponding top BLASTP hits in other eukaryotes, apicomplexan parasites, *Saccharomyces cerevisiae*, or *Homo sapiens*. Cyan stars represent annotated histone H3 genes. (B) Protein sequence alignment of H3 (cyan) and CENH3 (black) in *C. parvum*. Amino acids that are identical between at least two sequences are highlighted in grey. (C) Diagram of the targeting constructs designed to replace the endogenous *tk* locus with a second copy of either of the genes annotated as Histone H3-like proteins in *C. parvum* (*cgd3_2540* or *cgd4_3220*), a 3HA tag, and a Nluc-P2A-NeoR cassette. (D) Immunofluorescence staining of transgenic H3.1-3HA or (E) H3.2-3HA parasites. HCT-8 cells were infected with transgenic oocysts, fixed at 18 hpi, and stained with rat anti-HA followed by the secondary antibody Alexa Fluor 488 goat anti-rat IgG. Hoechst was used for nuclear staining. Images were acquired as Z-stacks using a Zeiss LSM-880 Laser Scanning Confocal microscope equipped with Airyscan (LSCM-A) and are presented with orthogonal views. Scale bars, 1 μm. Data represent two biological replicates, each with 3-5 technical replicates.

**Table 1:** The results of BLASTP analyses using the *C. parvum* centromeric histone H3-like protein (*cgd4_2030*), histone H3-like proteins (*cgd3_2540* and *cgd4_3220)*.

| Query | Top hits in Apicomplexa<sup>*</sup> | E value | Top hits in all Eukaryotes<sup>#</sup> | E value | Top hits in *Saccharomyces cerevisiae* | E value | Top hits in *Homo sapiens* | E value<sup>117</sup> |
| --- | --- | --- | --- | --- | --- | --- | --- | --- |
| *cgd4_2030* (CENH3) | histone H3-like centromeric protein CSE4 [*C. cionae*] | 5e-46 | centromeric DNA-binding histone H3-like protein cse4 [*N. ribis*] | 3e-39 | Chain A, Cse4 | 2e-30 | Chain A, Histone H3.1, Histone H3-like centromeric protein A, Histone H3.1 | 5e-39 |
| *cgd3_2540* (H3.1) | histone H3 [*C. cionae*] | 2e-86 | histone H3-like [*P. hypophthalmus*] | 3e-86 | histone H3 | 4e-81 | histone H3.2 | 1e-88 |
| *cgd4_3220* (H3.2) | histone H3 [*C. cionae*] | 2e-90 | histone H3-like [*P. hypophthalmus*] | 2e-87 | histone H3 | 2e-81 | histone H3.2 | 1e-89 |

*parvum* proteins were compared against all organisms in the Apicomplexa phylum (except for *Cryptosporidium* spp.),

**Table 2:** The identity and similarity results from multiple global sequence alignments between H3.1, H3.2, and CENH3 proteins using a BLOSUM62 scoring matrix.

| *Cp* H3.1 and *Cp* H3.2 | *Cp* CENH3 and *Cp* H3.1 | *Cp* CENH3 and *Cp* H3.2 |
| --- | --- | --- |
|  | Identity: 46.31%<br>Similarity: 65.10%<br>Gaps: 14.09% |  |
| Identity: 97.78%<br>Similarity: 99.26%<br>Gaps: 0.00% |  | Identity: 46.31%<br>Similarity: 64.43%<br>Gaps: 14.09% |

To further explore the distribution of H3.1 and H3.2 H3 variants, we generated two transgenic lines of parasites that replaced the dispensable thymidine kinase (*tk*) gene with a second copy of *cgd3_2540* (referred to as H3.1-3HA) or *cgd4_3220* (referred to as H3.2-3HA) fused with a triple hemagglutinin (3HA) epitope tag using CRISPR/Cas9 (**Fig 1C, S1 A-B Fig, Table S1**). Transgenic parasites were used to infect a human adenocarcinoma cell line (HCT-8) monolayer, cultures were fixed at 18 hours post infection (hpi) during merogony, stained, and the localization of the 3HA tag was observed by immunofluorescence (IFA) high-resolution confocal laser scanning microscopy equipped with Airyscan detection (LSCM-A). We found that H3.1-3HA (**Fig 1D**) and H3.2-3HA (**Fig 1E**) had a diffuse staining pattern, which was expected based on histone H3 staining in other apicomplexan parasites [27, 44, 45]

### The staining pattern of C. parvum centromeric histone H3 (CENH3) is distinct

To visualize *C. parvum* centromeres, we used CRISPR-Cas9 gene editing to add a 3HA tag to the C-terminus of the centromere-specific histone H3 variant protein (referred to as CENH3-3HA) (**Fig 2A, S3 A-B Fig, Table S1**). After successfully selecting and amplifying these transgenic parasites in immunocompromised mice, we infected HCT-8 cells for 22 h and examined the staining pattern by LSCM-A microscopy. For these studies, we chose mature meronts that were oriented with the basal side of the vacuole at the surface of the host cell where it attaches at the feeder organelle and where the majority of meronts were projecting upward, as ascertained by fluorescence staining and Z-stack images running perpendicular from bottom to top of the parasite (see cartoon on **Fig 3C**). The CENH3 staining pattern observed in *C. parvum* was unexpectedly highly diffuse and occupied the apical half of the nucleus (**Fig 2B-C**) in contrast to the more dispersed pattern found in the H3.1-3HA and H3.2-3HA tagged parasites (**Fig 1 D,E**). The diffuse pattern of CENH3 staining in *C. parvum* contrasts sharply with the focal pattern of regional centromeres previously described in *T. gondii* [27, 46]. Hence, we utilized a CENH3-3HA-tagged *T. gondii*, gifted by Dr. Kami Kim, as a staining control for our experiments. These parasites were used for the infection of HCT-8 cells, fixed at 24 hpi, stained, and examined by high-resolution LSCM-A immunofluorescence (IFA) microscopy (**Fig 2D-E)**. We found that the staining pattern of the centromeric histone H3 in *T. gondii* is punctate, similar to what has been found previously in *T. gondii* by other investigators [27, 46].

**Figure 2.**
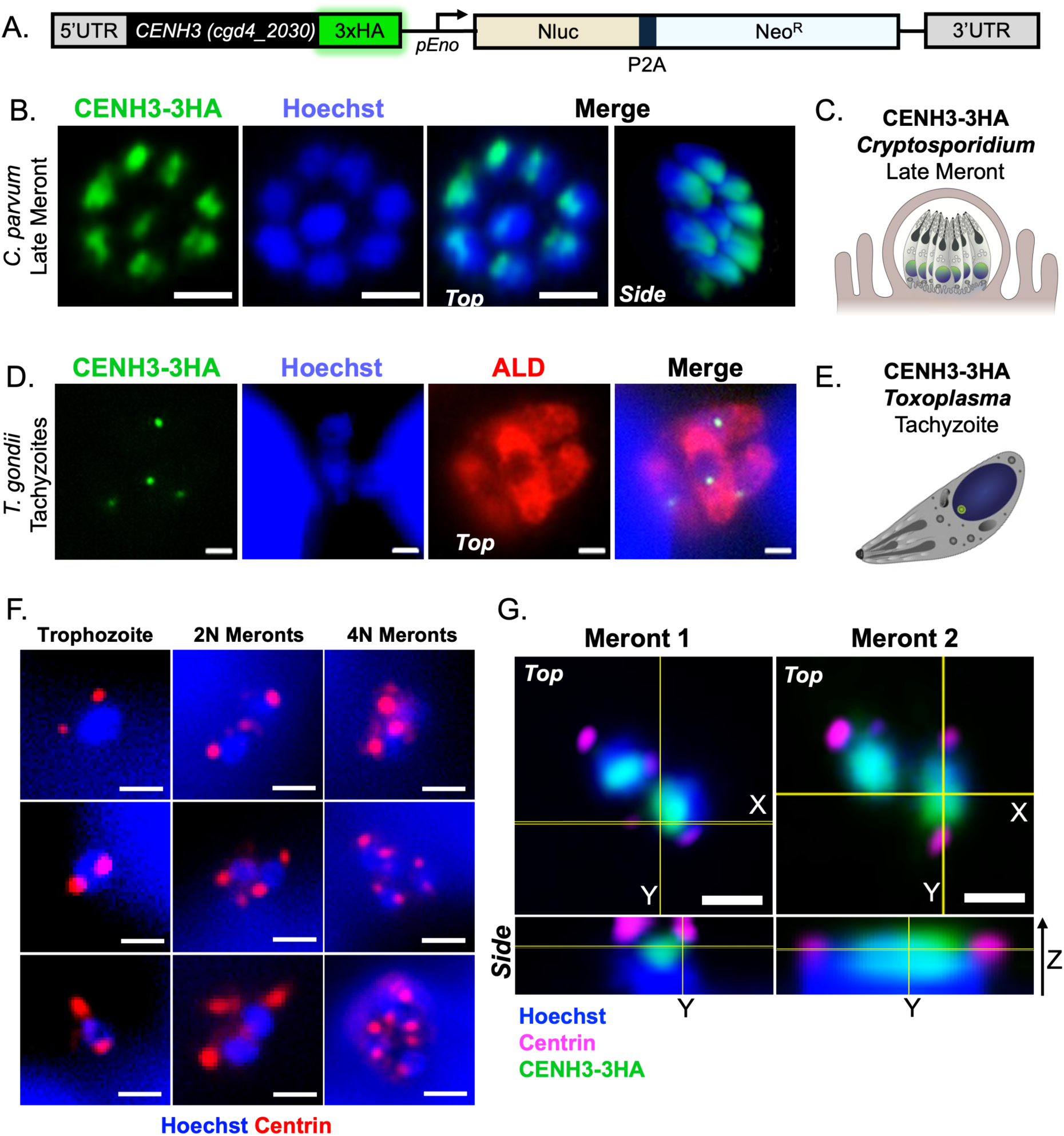
The staining pattern of *C. parvum* centromeric histone H3 (CENH3) is distinct from *Toxoplasma* and remains diffuse during mitosis. (A) Diagram of the targeting construct designed to add a 3HA tag and Nluc-P2A-Neo^R^ cassette to the C-terminus of the *C. parvum* CENH3 (*cgd4_2030*). (B) HCT-8 cells were infected with CENH3-3HA *C. parvum* parasites, fixed at 22 hpi, and stained with rat anti-HA, followed by the secondary antibody Alexa Fluor 488 goat anti-rat IgG followed by Hoechst staining. Images were acquired as Z-stacks using LSCM-A and are presented with orthogonal views. Scale bars, 1 μm. (C) Cartoon depiction corresponding to the microscopy image in panel (B). In *Cryptosporidium*, a late-stage meront contains eight nuclei that have recently completed cytokinesis to form mature merozoites. The CENH3-3HA signal within each nucleus displays a diffuse staining pattern in *C. parvum*. (D) HCT-8 cells were infected with CENH3-3HA *T. gondii* parasites, fixed at 24 hpi, and stained with rat anti-HA and rabbit anti-aldolase (ALD) to visualize the cytosol, followed by secondary antibodies Alexa Fluor 488 goat anti-rat IgG and Alexa Fluor 568 anti-rabbit IgG. Hoechst was used for nuclear staining. Images were acquired as Z-stacks using LSCM-A and are presented with orthogonal views. Scale bars, 1 μm. (E) Cartoon depiction corresponding to the microscopy image in panel (D). In *T. gondii* tachyzoites, the CENH3-3HA signal within each nucleus displays a discrete staining pattern. (F) HCT-8 cells were infected with excysted *C. parvum* sporozoites for 2 h, then washed twice to remove extracellular parasites. Cultures were fixed at 30 min increments between 6-9 hpi to identify the first (trophozoites with one nucleus), second (early meronts with 2 nuclei), and third (mid-stage meronts with 4 nuclei) mitotic divisions during the first round of merogony by widefield microscopy. Scale bars, 1 μm. (G) Cultures were infected with CENH3-3HA *C. parvum* sporozoites for 2 h, washed, and fixed at 6.5 hpi to capture the first round of mitosis. Cultures were stained with rat anti-HA, rabbit anti-centrin-1, followed by secondary antibodies Alexa Fluor 488 goat anti-rat IgG, Alexa Fluor 568 goat anti-rabbit IgG, and lastly Hoechst. Parasites undergoing mitosis were identified by having two centrin-1 points per nucleus, whereas parasites in interphase had only one centrin-1 point. Images were acquired as Z-stacks using LSCM-A and are presented with orthogonal views. Scale bars, 1 μm. Data represent 2-3 biological replicates, each with 3-5 technical replicates.

**Figure 3:**
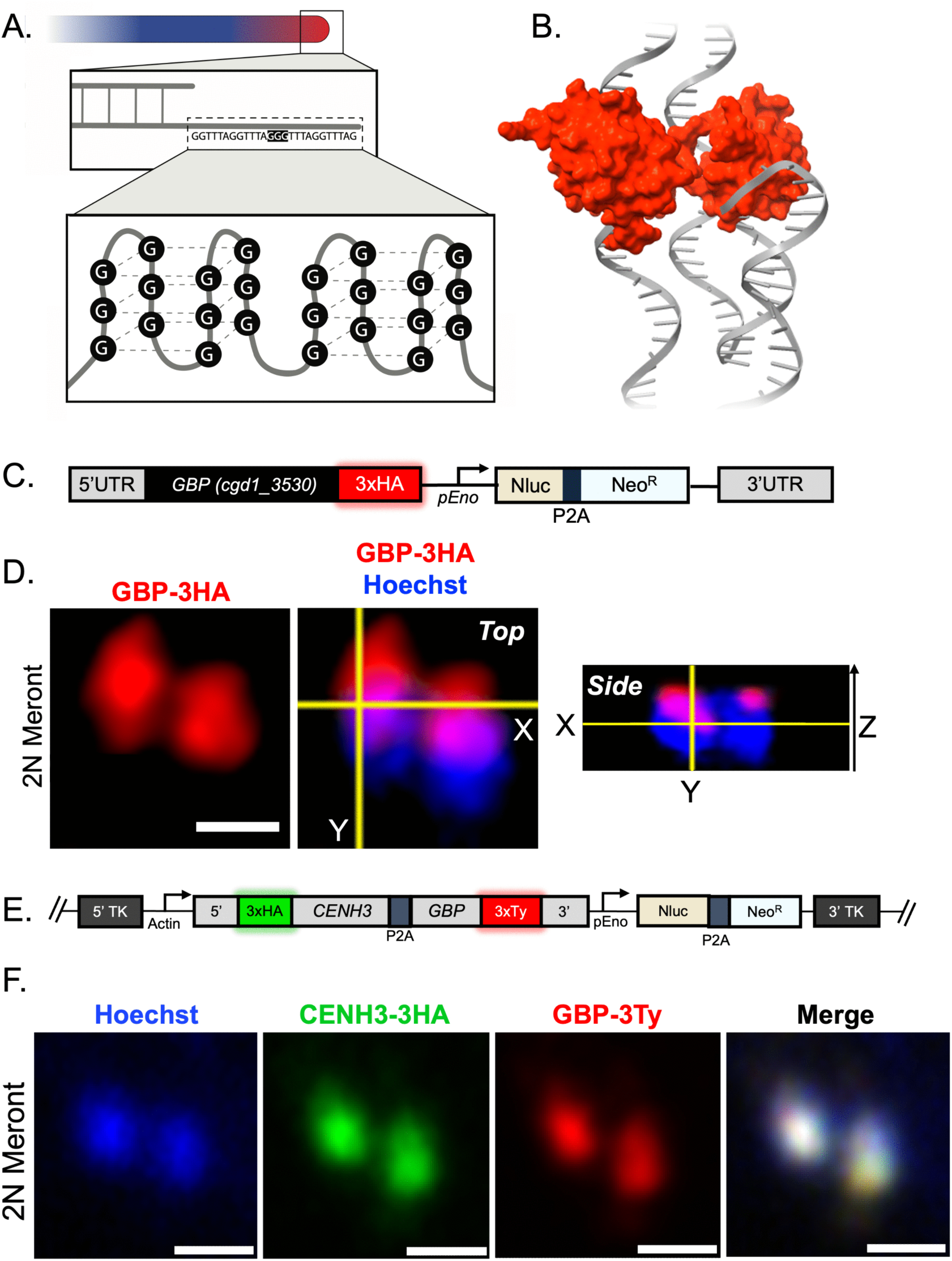
The telomere (G-strand binding protein, GBP) staining pattern is apical in *C. parvum* and overlaps with the centromere (CENH3) staining pattern. (A) The 3’ overhang single-stranded DNA at the end of telomeres contains repetitive sequences (GGTTTA)_n_ in *C. parvum* that form G-quadruplex structures. (B) The G-quadruplex binding protein (GBP) is essential for forming and stabilizing these structures. The sequence of *Cp*GBP and its binding motif has been previously determined and the protein structure is modeled by Alphafold and ChimeraX [51]. (C) Diagram of the targeting construct designed to add a 3HA tag and Nluc-P2A-Neo^R^ cassette to the C-terminus of the G-quadruplex binding proteins gene (GBP*, cgd1_3530)* (D) HCT-8 cells were infected with GBP-3HA parasites, fixed at 6.5 hpi to capture early meronts (2 nuclei), and stained with rat anti-HA, followed by the secondary antibody Alexa Fluor 488 goat anti-rat IgG. Hoechst was used for nuclear staining. Images were acquired as Z-stacks using LSCM-A and are presented with orthogonal views and the scale bar is 1 μm. (E) Diagram of the targeting construct designed to replace the endogenous *tk* locus with a second copy of CENH3 N-terminally tagged with 3HA and GBP with a C-terminal 3xTy tag separated with a P2A split peptide. (F) Immunofluorescence staining of transgenic 3HA-CENH3-GBP-3Ty parasites. HCT-8 cells were infected with transgenic oocysts, fixed at 6.5 hpi to capture early meronts (2 nuclei), and stained with rat anti-HA and mouse anti-Ty, followed by a secondary antibody stain containing Alexa Fluor 488 goat anti-rat IgG and Alexa Fluor 568 goat-anti mouse. Hoechst was used for nuclear staining. Images were acquired as Z-stacks using LSCM-A and are presented with orthogonal views. Scale bars, 1 μm. Data represent 2-3 biological replicates, each with 3-5 technical replicates.

### Distribution of centrosomes in C. parvum

A vital step in mitosis is centrosome duplication, and previous studies in apicomplexans have shown that it can be traced using antibodies that recognize the centrin 1 protein [27, 47]. A single point adjacent to the nucleus occurs in non-replicating cells, while two centrin 1 positive points per nucleus indicates the parasite is undergoing replication [27, 47]. In *Plasmodium*, during schizogony, asynchronous duplications of centriolar plaques, centrosomes, have been observed across multiple nuclei within a single cytoplasm, in this study we sought to identify analogous centrosomes in *C. parvum*. [28]. To identify parasites undergoing early mitosis, we infected HCT-8 cells with excysted wild type *C. parvum* sporozoites, washed off uninvaded sporozoites at 2 h, fixed the cultures at various timepoints between 6.5-8.5 hpi, and stained with Hoechst and an anti-centrin-1 antibody. Initially, trophozoites contained a single centrin-1 focus associated with the single nucleus (**S4A**). However, in other examples of trophozoites, we often observed that each nucleus consistently exhibited two centrin-1 positive points (**Fig 2F**). Furthermore, by visualizing the number of nuclei based on Hoechst staining, and centrin-1 to monitor dividing centrosomes, we observed that all nuclei found in a common cytoplasm shared the same centrin-1 staining pattern. For example, when 2N or 4N meronts were observed to have duplicated the centriole as shown by centrin-1 staining, the duplicated pattern was consistently observed for all nuclei within that cell (**Fig 2F**). To determine if the diffuse centromere staining pattern persisted during active mitosis in *C. parvum*, we infected cultures with CENH3-3HA parasites and repeated the experiment previously described. We found that nuclei undergoing mitosis also displayed a diffuse CENH3-3HA staining pattern by LSCM-A (**Fig 2G**). We also attempted to localize CENH3 using ultrastructure expansion or immuno-EM staining methods. Despite extensive troubleshooting, we could not identify positive signals in N– or C-terminally tagged CENH3-3HA lines, or in separate line expressing a spaghetti monster (sm)HA tag (i.e. CENH3-smHA), by either method, which may be due to the low abundance of CENH3 or difficulty preserving the epitope during these procedures.

### The telomere (G-strand binding protein, GBP) staining pattern is apical in C. parvum

Previous studies in *Plasmodium* have found that the telomeres are consistently found on the opposite pole to the centromeres [48–50]. Since the centromeres were located toward the apical end of the nuclei in meronts, we expected that *C. parvum* telomeres would be more basal. To examine the distribution of telomere in *C. parvum*, we utilized a surrogate protein marker. G-quadruplex structures readily form at repetitive telomeric sequences and require G-quadruplex binding proteins (GBPs) for stabilization (**Fig 3A**). A previous study identified *cgd1_3530* as the GBP in *C. parvum* and determined the specific sequence necessary for *Cp*GBP binding (GGTTA) [51]. Using AlphaFold and ChimeraX, we modeled the G-quadruplex structure formed by this repetitive sequence and visualized how the *Cp*GBP protein interacts with and stabilizes it. (**Fig 3B**). To identify the location of *C. parvum* telomeres relative to the centromeres, we engineered transgenic parasites to express a 3HA tag at the gene encoding the C-terminus of the G-quadruplex binding protein (referred to as GBP-3HA) (**Fig 3C, S3 A-B Fig, Table S1**). LSCM-A microscopy on infected HCT-8 monolayers (24 hpi) revealed that the telomeres were located toward the apical side of the nucleus (**Fig 3D**), rather than at the opposite pole as expected.

### The telomere (GBP) and centromere (CENH3) staining patterns overlap in C. parvum

To confirm that *C. parvum* centromeres and telomeres had overlapping nuclear locations, we generated an additional line of transgenic parasites that replaced the dispensable thymidine kinase (*tk*) gene with a second copy of an N-terminally 3HA tagged CENH3 and GBP C-terminally tagged with 3Ty via CRISPR/Cas9, with an intervening P2A split peptide to facilitate co-expression of separate proteins (referred to as 3HA-CENH3-GBP-Ty) (**Fig 3E, S1C-D Fig, Table S1)**. HCT-8 cultures were infected with 3HA-CENH3-GBP-Ty parasites, fixed at 6.5 hpi, and examined by LSCM-A microscopy after staining. We observed that 3HA-CENH3 and GBP-3Ty signals were both more concentrated in the apical region of the nucleus, as defined by Hoechst staining. This overlap was evident by fluorescence microscopy and further supported quantitatively using the Coloc2 plugin in FIJI/ImageJ, which yielded a Pearson correlation coefficient of 0.96 (**Fig 3F, S2 Fig)**.

### Detection of mitotically active DNA, microtubules, and the nuclear pore proteins in C. parvum

The phosphorylation of histone H3 (specifically at serine 10 and serine 28) is a highly conserved mechanism for initiating chromosome condensation in preparation for mitosis [52]. Antibodies recognizing the phosphorylated version of H3 at serine 10 are commonly used in other systems to determine which cells are currently undergoing mitosis from late interphase through telophase [53]. Even though apicomplexan parasites are known not to condense their chromosomes, *C. parvum* meronts frequently stained positive for the phosphorylated histone H3 (PH3) antibody (**Fig 4A**), suggesting they are in the process of mitosis. Previous studies have characterized the distribution of nuclear pores in *Plasmodium falciparum* using a rabbit polyclonal antibody to the protein NUP116 [54]. We rationalized that the epitope recognized by this antibody might also be conserved in *C. parvum*, despite their evolutionary distance. We visualized nuclear pores in *C. parvum* using the rabbit anti-NUP116 antibody and found that it reliably stained the parasite nucleus in a punctate pattern that was concentrated primarily peripherally in the nucleus (**Fig 4B**). Although we have not verified the identity of this cross-reaction, it presumably represents a component of nuclear pores that will be used for future studies on nuclear organization in *C. parvum*.

**Figure 4:**
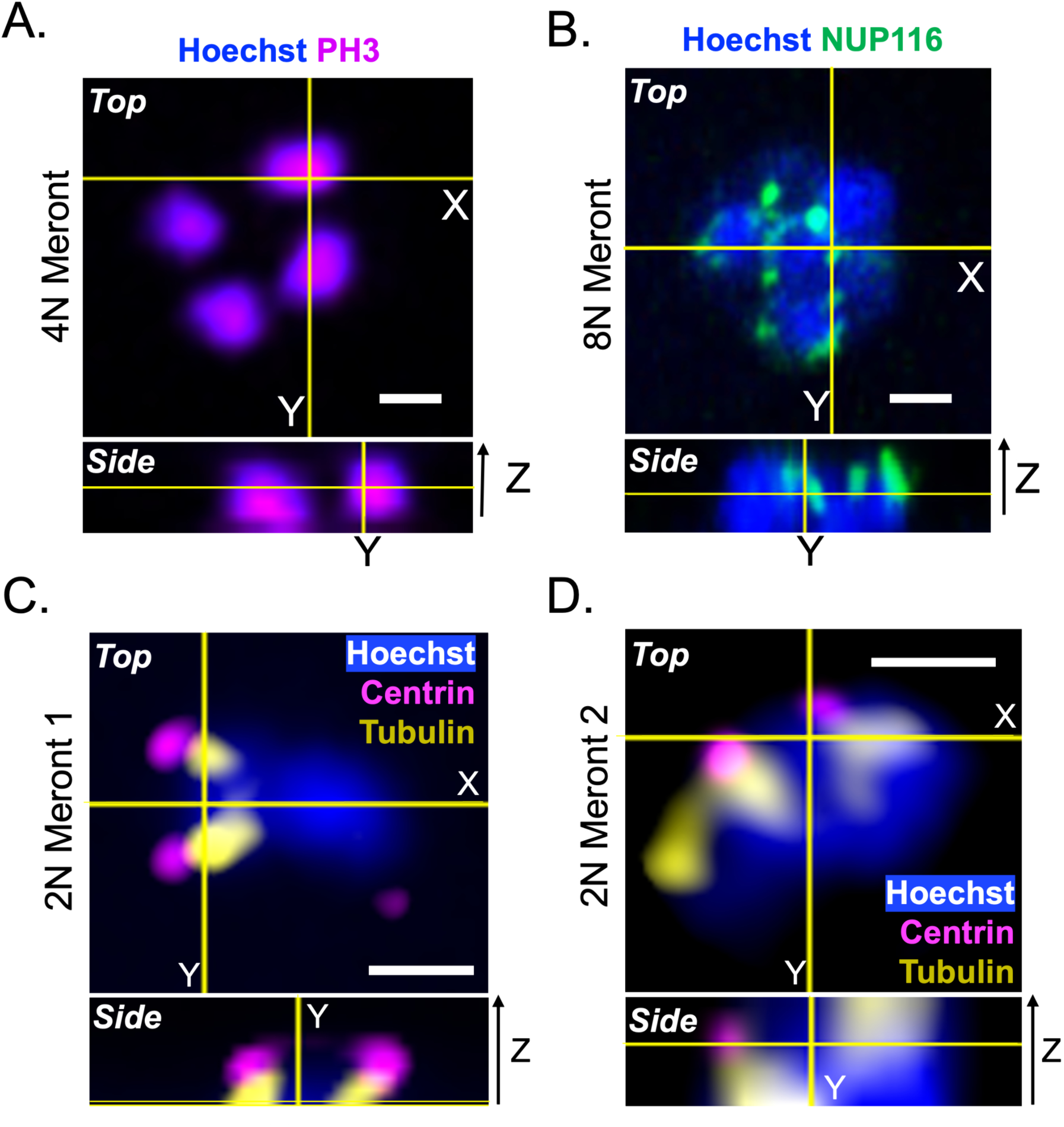
Mitotically active DNA, microtubules, and the nuclear pore proteins as visualized by antibody staining in *C. parvum*. (A) HCT-8 cells were infected with wild type parasites, fixed at 18 hpi, and stained with rabbit anti-phosphorylated Histone H3 (PH3) followed by an Alexa Fluor 568 goat anti-rabbit IgG antibody. DNA was stained with Hoechst and a mid-stage meront with 4 nuclei is shown. (B) Parasites were prepared identically to panel A, but were stained with a rabbit polyclonal antibody that recognizes a subunit of the nuclear pore in *Plasmodium falciparum* (NUP116) followed by an Alexa Fluor 568 goat anti-rabbit IgG antibody and Hoechst. A late-stage meronts with 8 nuclei is shown. (C, D) HCT-8 cells were infected with wild-type parasites, fixed at 6.5 hpi to capture early meronts (2 nuclei), and stained with rabbit anti-centrin-1 and mouse anti-tubulin, followed by the secondary antibodies Alexa Fluor 488 goat anti-mouse IgG, Alexa Fluor 568 goat anti-rabbit IgG, and Hoechst. The nucleus in the first meront (C) may be an example of early mitosis, whereas the second meront (D) may be an example of a later phase of the mitotic process. Images were acquired as Z-stacks using LSCM-A and are presented with orthogonal views. Scale bars, 1 μm. Data represent two biological replicates, each with 2-5 technical replicates.

Among apicomplexan parasites, mitosis has primarily been studied in *T. gondii* and *P. falciparum* [16, 17, 27, 55]. These prior studies demonstrate that once centrosomes duplicate, and the nuclear envelope forms a tunnel-like structure, organizing spindle microtubules within it, spindle microtubules then attach at the centromere and mediate chromosome movement [56]. We also attempted to visualize this process in *C. parvum* by staining with antibodies that recognize the centrin-1 nd alpha tubulin (**Fig 4C-D, S4B-C**). We examined cells at 6-7 hpi to capture early stages of mitosis. We frequently observed two conformations of centrin-1 tubulin structures that may represent early (**Fig 4C, S4B**) and late mitotic phases (**Fig 4D, S4C**). In Fig 4C, the duplicated centrin signal shows a single discrete spindle (microtubule signal) overlying what appears to be a single nucleus, as evident by size and shape. In other examples the centrin spots have moved further away, and the spindles elongated (Y-shaped) (**Fig 4D, S4C**). Notably, the nucleus in this image is much larger and diffuse, consistent with it being at a later stage that is starting to divide.

### Composition of centromeres in C. parvum and T. gondii

Centromeres are defined by the deposition of CENH3-containing nucleosomes [27, 57]. Given the diffuse staining of CENH3, we wanted to further characterize the centromere by identifying DNA sites bound by this protein. Therefore, we adopted protocols for CUT&RUN to identify centromeres based on CENH3 binding based on capture with anti-HA antibody in *C. parvum* [43]. To ensure sufficient HA binding, we developed a CENH3 line tagged with <u>s</u>paghetti <u>m</u>onster (referred to as CENH3-smHA) that contains 10 HA epitope repeats for these experiments and validated identical staining patterns to CENH3-3HA *C. parvum* (**S3 A-C Fig, Table S1**). To validate our experimental strategy, we first applied CUT&RUN to *T. gondii*, whose centromeres have previously been defined by ChIP-seq [27]. Our CUT&RUN data closely matched earlier findings, with CENH3 enrichment observed at discrete chromosomal loci corresponding to the previously described centromeres (**Fig 5A, S5A-C Fig**). We also detected the positive correlation of CENH3 with H3K9me3 and inverse correlation with H3K4me3 reported in earlier ChIP-chip analyses (**Fig 5A, S5A-B Fig**). Based on these signals, we implemented a stepwise strategy to identify centromeric regions. Valleys in H3K4me3, consisting of 20 kb windows, sized to match reported centromere lengths (16 kb ± 3.4 kb)[27], combined with co-enrichment of H3K9me3 with CENH3-smHA were used as criteria for identifying centromere regions (**S5D Fig**). Applying these rules, we identified a single centromere in each of the 13 chromosomes of *T. gondii* (**S5E Fig**), with Chr VIIb identified as part of Chr VIII [58](**S5C Fig**).

**Figure 5.**
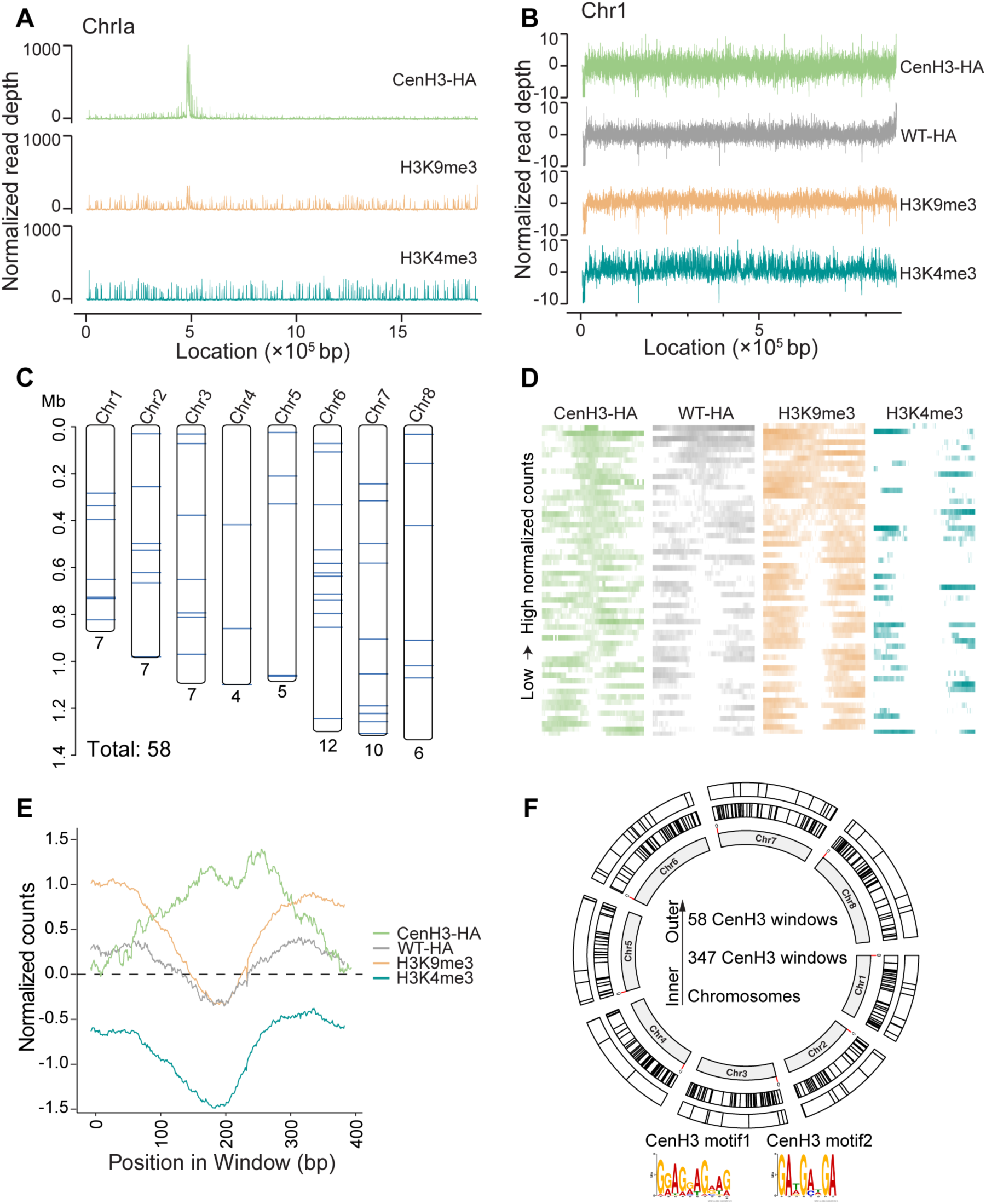
Genome-wide distribution of CENH3 in *Toxoplasma gondii* and *Cryptosporidium parvum*. (A) Genome browser view of *T. gondii* chromosome Ia showing CUT&RUN signals for CENH3-HA (HA antibody), H3K9me3, and H3K4me3. CENH3 signal enrichment positively correlates with H3K9me3 and inversely with H3K4me3. Peaks of CENH3 enrichment are indicated (red arrows). (B) Genome browser view of *C. parvum* chromosome I, showing CUT&RUN signals for CENH3-HA *C. parvum* captured with HA antibody and wild type *C. parvum* captured with HA antibody (referred to as WT-HA), H3K9me3, and H3K4me3 (captured with antibodies). (C) Chromosomal distribution of the 58 CENH3-enriched windows in intergenic regions of *C. parvum* chromosomes (Chr#). Numbers below each chromosome indicate the number of windows. (D) Heatmaps of CUT&RUN signals for the groups shown in (B) within a 400 bp window around all 58 CENH3-enriched windows. Each row represents an individual centromeric site, ranked by signal intensity. (E) Average signal profiles for CENH3-HA (green), WT-HA (gray), H3K9me3 (orange), and H3K4me3 (teal) across the 400 bp window around CENH3 sites. (F) Circos plot of CENH3-associated motifs. From inner to outer rings: chromosomes, locations of 347 CENH3-enriched windows found in coding regions (black), and locations of 58 CENH3-enriched windows found in intergenic regions (black). Representative sequence logos for motif 1 and motif 2 are shown.

We next applied CUT&RUN to *C. parvum*. Initial experiments using a 120-minute MNase digestion failed to generate usable data, as both sequencing depth and genome coverage were low compared to negative controls (**S1 Dataset**). Since CENH3 nucleosomes can be hypersensitive to MNase digestion, as reported previously in *Caenorhabditis elegans* (*C. elegans)* [57], we tested digestion times of 5, 30, and 60 min. The 30 min digestion yielded the best coverage and sequencing depth, and these conditions were used for following analyses (**S1 Dataset, S6A Fig**). Notably, sequencing depth of the *C. parvum* samples was more variable and, in some cases, lower than that of *T. gondii*. This difference primarily reflects technical challenges associated with generating transgenic *C. parvum* parasites. To ensure data quality and comparability, we included only samples with high genome coverage (>80%) in downstream analyses. Unlike *T. gondii*, read mapping profiles for CENH3 in *C. parvum* showed signals broadly distributed along chromosomes (**Fig 5B**). To locate centromeric regions in *C. parvum*, we first attempted the *T. gondii* stepwise strategy, but no valid windows were detected. As the CENH3 distribution resembled that of the holocentric nematode *C. elegans*, we adjusted thresholds accordingly by restricting the peaks analysis to 400 bp windows (**S6B Fig**). We identified 423 CENH3-enriched regions distributed across all chromosomes. Heatmaps and aggregate signal profiles showed that these windows displayed features of CENH3 enrichment correlated with H3K9me3 deposition and H3K4me3 depletion, suggesting that they represent centromere-associated chromatin (**S6C-D Fig**). Importantly, these signals were largely absent in wild type parasites stained with HA antibody as a control, supporting their authenticity (**S6C-D Fig**). To further refine localization, we excluded windows overlapping with WT-HA signals, reducing the dataset to 347 high-confidence centromeric windows (**S6B,E Fig**). Taken together, these results indicate that *C. parvum* possesses many small CENH3 binding sites dispersed across eight chromosomes, in contrast to the monocentric nature of *T. gondii*.

We further investigated characteristics of *C. parvum* centromeres. In contrast to *T. gondii*, where centromeres are typically located within large intergenic regions [27], no extended intergenic regions were observed in *C. parvum* due to the tight compaction of the genome. Surprisingly, the majority of *C. parvum* CENH3-associated windows were located within gene coding regions rather than being restricted to intergenic regions (**S2 DataSet2**). To evaluate evolutionary constraints on CENH3-associated regions, we performed nucleotide diversity (Pi) analyses on the DNA sequences within these regions and dN/dS analyses on the genes overlapping these windows. We compared previously generated genomic sequences from numerous *C. parvum* IIa and IId isolates, which show high genomic similarity. Most CENH3-associated windows (∼82%) showed reduced Pi values compared with genome-wide 400-bp sliding windows, and only a single gene within the windows exhibited an elevated dN/dS ratio relative to the genome-wide orthologous gene set (**S6F,G Fig**). These results indicate that CENH3-associated regions are highly conserved and under strong purifying selection. To determine whether centromeric regions were also present within intergenic regions of *C. parvum*, we applied a stepwise identification strategy using a slightly relaxed read depth threshold (**S6F Fig**). We identified 58 centromeric regions distributed within intergenic regions across all chromosomes (**Fig. 5C, S3 DataSet**). Heatmaps and aggregate signal profiles for CNEH3 as well as chromatin marks also supported that they represent centromere-associated chromatin (**Fig. 5D,E**). These results suggest that CENH3 binding in *C. parvum* is broadly distributed throughout the chromosomes, including both with in coding regions and intergenic regions. To assess sequence properties of *C. parvum* centromeric windows, motif analysis of *C. parvum* centromeric windows was performed using MEME, revealing two GA-rich motifs enriched at CENH3-associated sites (**Fig 5F**). Notable, these GA repeats are also found in other regions of the genome that were not identified in the CENH3 capture studies.

## DISCUSSION

Despite rapid asexual proliferation being required for parasite expansion, the molecular mechanisms that underlie *C. parvum* mitosis have remained unexplored. To investigate this process, we examined nuclear division using markers to track multiple rounds of mitosis during merogony. Nuclear division in *C. parvum* can be visualized by staining with phospho-H3 and centrin-1, which provide markers for mitotically active chromatin and duplicating centrosomes, respectively. Surprisingly, both CENH3, a marker for centromeres, and GBP, a marker for telomeres, were diffusely localized across the apical half of the nucleus. Moreover, these diffuse apical patterns persisted throughout mitosis, as shown by visualizing nuclear division stages with centrin-1 and microtubules. To explore the molecular nature of centromeres, we performed CUT&RUN sequencing of transgenic parasites expressing epitope-tagged CENH3. Although discrete monocentric peaks were detected in the related parasite *T. gondii*, CENH3 binding sites were diffusely scattered across ∼ >400 sites in the *C. parvum* genome. Overall, these findings indicate that *C. parvum* has evolved an alternative chromosome structure compared to other apicomplexan parasites, differences that may have significant functional consequences for chromosome stability and meiotic recombination.

Following entry of sporozoites (or merozoites) into the host cell, the parasite undergoes merogony, consisting of 3 rounds of nuclear division prior to cytokinesis to generate infectious merozoites. By visualizing the number of nuclei based on Hoechst staining, and centrin-1 to monitor dividing centrosomes, we observed that all nuclei found in a common cytoplasm shared the same centrin-1 staining pattern (single centrin-1 spot when not dividing and duplicated centrin-1 spot prior to nuclear division). This pattern suggests that nuclei within a single meront may divide synchronously, although further studies using synchronized cells and live cell imaging would be needed to substantiate this hypothesis. This pattern differs from *Plasmodium* schizogony, where nuclei are observed at different mitotic stages within a single schizont [28]. In *Plasmodium*, nuclear pores are important for division as they facilitate the transport of mitotic molecules through the intact nuclear envelope and associate with the centriolar plaques [31, 59]. Although we did not examine the dynamics of nuclear pores during division, they appear as peripheral puncta in mature meronts and may serve a similar function in *C. parvum*.

Staining with alpha-tubulin allowed us to visualize *C. parvum* microtubules during mitosis, particularly at early stages where microtubules form a cluster that may represents the spindle that will drive chromosome separation during mitosis. The pattern of microtubules described here in dividing cels is quite different from the stable, elongate microtubules that emanate from the apical pole of sporozoites and merozoites [60]. Such dramatic re-organization of microtubules between non-dividing and dividing cells is not unusual for eukaryotic cells, although the dynamics and regulation of this process have not been examined extensively in *Cryptosporidium*. Based on qualitative observations of centrosome duplication, centrin-1 positioning, microtubule organization, and changes in nuclear morphology, our studies suggest an approximate sequence of events corresponding to sequential rounds of mitosis during early, mid, and late merogony. Further studies using fluorescently tagged proteins and life cell imaging, combined with synchronous cell division would be important to validate and extend this model. Regardless of the exact dynamics of mitosis, these studies provide an important foundation for studies of the dynamics of centromere staining described below.

Unexpectedly, we found a diffuse nuclear CENH3-3HA staining pattern rather than the punctate pattern observed in other apicomplexan parasites [27, 29, 34]. We also confirmed that the diffuse CENH3 staining pattern did not deviate when the method of tagging was altered (e.g. smHA), when the location of the tag was swapped (N-terminally tagged, e.g. 3HA-CENH3-GBP-3Ty), or when a tagged second copy of the gene was inserted into a neutral locus of the genome (3HA-CENH3-GBP-3Ty). We also developed epitope-tagged lines of telomere-binding protein GBP, similar to studies in *Eimeria* and *Plasmodium* [61, 62]. Unlike in other apicomplexans, GBP was also localized to the apical half of the nucleus in *C. parvum*, overlapping closely with CENH3 [18–20, 48]. We further confirmed that the telomeres and centromeres occupied a similar nuclear location using a parasite strain in which CENH3 and GBP proteins were simultaneously expressed as tagged second copies. This pattern contrasts sharply with other apicomplexans where telomers are localized to the opposite pole of the nucleus from centromeres [48–50]. We sought to determine if centromeres remained diffuse throughout mitosis by simultaneously visualizing the centrosomes and centromeres by staining CENH3-3HA with anti-HA and centrin-1 antibodies. We confirmed that in *C. parvum,* the diffuse centromere staining pattern does not change throughout mitosis, which further contrasts with the highly focal nature of centromeres in *T. gondii* and *P. falciparum* [26, 27].

Given the highly unusual staining patterns of CENH3 in *C. parvum*, we decided to investigate CENH3 binding sites on DNA using CUT&RUN sequencing. For comparison, we also performed CUT&RUN studies on CENH3 in the related parasite *T. gondii* and compared them to previous studies that used ChIPSeq to define centromeres [27]. In *T. gondii*, CUT&RUN sequencing identified highly similar CENH3 binding sites to those reported previously by ChIPSeq [27] with a single predominant peak on each chromosome, including newly defined peaks on chromosome IV, that had not been identified previously. Centromeres in *T. gondii* consist of gene-poor regions of ∼15 kb that are not defined by any particular sequence or repeats but instead are positively correlated with H3K9me3 and inversely correlated with H3K4me3, typical chromatin patterns expected for centromeric regions [18, 27]. Under similar conditions, CUT&RUN experiments for CENH3 binding sites in *C. parvum* failed to identify similar singular dominant peaks on each chromosome. Instead, CENH3 binding sites were hypersensitive to MNase digestion, with the signal being the highest at a 30-min incubation time, similar to that observed in ChIP-seq studies of the holocentric nematode *Caenorhabditis elegans* (*C. elegans*) [57]. Under optimal digestive conditions, we found that the CENH3 signal in *C. parvum* was broadly distributed along chromosomes, with ∼ 350 distinct sites in coding regions and another ∼ 60 sites within intergenic regions scattered across 8 chromosomes. Interestingly, like *T. gondii*, *C. parvum* CENH3 peaks also correlated with H3K9me3 signal and were found in areas lacking H3K4me3, consistent with a centromere-associated chromatin pattern. The distribution of CENH3 binding sites in *C. parvum* is similar to that described in *C. elegans* as polycentric, implying that CENH3 binds to distinct sites rather than diffusely across the entire chromosome [57].

Collectively, our findings are consistent with *C. parvum* containing holocentric centromeres based on the following criteria: Firstly, the staining pattern of CENH3 diffusely stains the apical half of the nucleus under nondividing conditions, in contrast to the focal staining of CENH3 in *T. gondii* [18, 27] and *P. falciparum* [26]. Secondly, the diffuse pattern of CENH3 staining remains throughout mitosis, suggesting centromeres do not coalesce into a single regional center. Thirdly, CENH3 binding sites as defined by CUT&RUN resemble the dispersed pattern of low-intensity peaks seen in *C. elegans*, which has well-characterized holocentric chromosomes. Although we interpret this diffuse distribution as consistent with holocentric centromeres, we acknowledge that an alternative explanation, such as asynchronous chromatid separation or cohesion fatigue, could also contribute to the observed pattern. However, current imaging resolution does not allow simultaneous visualization of all microtubules and all CENH3-associated kinetochore sites, so we cannot fully distinguish between these possibilities. Therefore, we present our interpretation cautiously while noting that the number of predicted CENH3 loci substantially exceeds the 8 (1N) regional centromeres expected for a monocentric organization. Finally, the CENH3 binding sites show evidence of simple GA nucleotide repeats, a feature also seen in *C. elegan*s [57]. In both cases, the GA repeats are also found in other regions of the respective genomes, so this feature alone is not sufficient to mediate CENH3 binding. One unusual feature of the CENH3 binding sites in *C. parvum* is that they extend into coding regions, which is not an expected feature of centromeres. Interestingly, these regions were under strong purifying selection at the intraspecies level. The dispersed polycentric pattern of CENH3 binding sites in *C. elegans* is also unusual in that these regions are concentrated in High Occupancy Targets (HOT) sites for permissive binding of transcription factors [57]. In *C. elegans*, the CENH3 and HOT sites appear to serve a dual purpose: in replicating cells they bind CENH3, while they recruit transcription factors when CENH3 is lost during exit from the cell cycle in differentiated muscle cells [57]. By analogy, transcriptional silencing during mitosis in *C. parvum* may allow CENH3 binding in coding regions to facilitate chromatid segregation, with displacement during the rest of the cell cycle when transcription is active. Alternatively, the CENH3 binding sites in intergenic regions may function as the primary centromere-localizing signal for formation of kinetochores.

Holocentric chromosomes have been documented in many species, primarily through direct cytological observation, yet their functional implications and evolutionary origins remain poorly understood [37, 57, 63, 64]. Recent comparative and modeling work in fungi shows that complete transitions between centromere ‘types’ (point/regional/holocentric) occur progressively and do not fix by drift alone; rather, selection is needed to drive and stabilize new architectures while maintaining compatibility with the relatively conserved kinetochore–spindle interface [65]. In this light, *the* many small CENH3-binding sites in *C. parvum* fits a model of progressive centromere remodeling along an evolutionary continuum, rather than representing an abrupt departure from other apicomplexan centromere types.

The organization we describe has clear functional implications, as the presence of hundreds of small CENH3-positive sites suggests that *C. parvum* chromosomes may rely on numerous ‘micro-kinetochores’ distributed along their length. By spreading spindle attachment across many small units rather than concentrating mechanical force at a single centromere, the parasite could effectively diffuse the load generated during chromosome segregation. Such a system would be particularly advantageous for *C. parvum*, which performs closed mitosis with uncondensed chromatin and must execute multiple rounds of synchronous division during rapid merogony. In this framework, the diffuse, apically clustered distribution of CENH3 is not simply a structural anomaly, but an evolved solution to the mechanical and cell-cycle constraints imposed by rapid, multi-round division in an intestinal environment. Thus, the centromere architecture of *C. parvum* likely reflects selective pressures associated with maintaining chromosome stability, rather than a simple deviation from typical apicomplexan chromosome biology.

Current evidence suggests that holocentric chromosomes have evolved convergently at least 13 times across distantly related taxa in both the plant and animal kingdoms [36, 37, 64]. Holocentricity has not previously been described in protists, and hence it likely evolved convergently in *C*. *parvum*. However, we note that the directionality of centromere evolution within Apicomplexa remains unresolved. Because centromere organization has not been characterized in closely related outgroups such as gregarines or chromerids, it is equally plausible that the common ancestor of Apicomplexa possessed holocentric chromosomes and that monocentric organization arose secondarily in coccidia and hematozoa. Thus, the holocentric architecture in *C. parvum* may represent either an independently derived state or the retention of an ancestral condition. It is also possible that holocentricity is more widespread among protists, but that it has been overlooked due to their small size and failure to condense chromosomes during division.

One proposed advantage of holocentricity is its capacity to stabilize chromosomal fragments during mitosis, thereby mitigating genetic loss. This trait may confer a selective benefit to organisms frequently exposed to elevated levels of DNA damage due to environmental stressors or lifestyle factors [36, 63]. Thus, holocentricity may impart genome stability under stress encountered in the gastrointestinal system where the parasite is exposed to toxic bacterial metabolites [66] and host immune responses [67]. Despite this potential advantage, holocentricity also presents a major challenge for coordinating chromatid segregation following crossover during meiosis I. Well-characterized holocentric species have evolved mechanisms to circumvent this issue, such as inverted meiosis or, in the case of *C. elegans*, the deployment of kinetochore proteins that localize to discrete chromosomal regions, effectively mimicking monocentric behavior [68]. Whether *C. parvum* uses a similar mechanism as *C. elegans* to localize kinetochore proteins to discrete sites on the chromosome is uncertain, but previous work from our laboratory examining crossover rates during *C. parvum* meiosis revealed a high frequency of crossover events and failed to find evidence for inverted meiosis, a process used by some holocentric organisms [69]. In contrast, interspecies crosses between *C. tyzzeri* and *C. parvum* revealed that crossovers are strongly suppressed and most chromosomes are inherited uniparentally without crossovers [70]. Although we have not interrogated the nature of CENH3 binding sites in *C. tyzzeri*, differences in the recognition sequences or pattern of centromere distribution (i.e. monocentric vs. holocentric) could result in suppression of hybrids due to miotic or meiotic interference.

### Limitations and Future Studies

Examination of the nuclear organization of *C. parvum* throughout the cell cycle revealed a persistently diffuse centromere staining pattern, suggesting a holocentric chromosome structure. During division, such multiple centromeric sites dispersed along the chromosome are expected to bind centrosomes and thereby connect to microtubules for chromosome separation. Although we visualized events consistent with this process in an artificial timeline, future studies using fluorescent reporters could potentially provide more precise information on the formation of centrosomes, coordinated assembly of microtubules, and nuclear division in *C. parvum*. However, obtaining direct cytological data in support of holocentric centromeres and their behavior during nuclear division will be challenging in this system because chromosomes do not condense during division. Alternatively, it may be possible to explore the nature of centromeres by examining the distribution of other conserved centrosomes and kinetochore binding proteins during nuclear division. Such studies could shed further light on chromosome tethering in nuclear division and provide invaluable insights into unique evolutionary adaptations in *C. parvum*. Additionally, extending these studies to define how centromeres are organized in related species could provide insight into mechanisms of diversification that drive speciation.

## Acknowledgements

We thank Jennifer Barks for assistance with cell culture, Dr. Wandy L. Beatty, Microbiology Imaging Facility, Washington University School of Medicine in St. Louis, for assistance with microscopy, and Dr. Kami Kim, Univ. South Florida, for providing the *T. gondii* CENH3-HA line used here. We are grateful for helpful comments from Drs. Daniel E Goldberg, Douglas L. Chalker, and Tim Schedl, Washington University and Dr. Lihua Xiao, South China Agricultural University. Genome sequencing was performed by the GTAC Genome core at Washington University in St Louis. Supported in part by the Graduate Research Fellowship Program, National Science Foundation (to A.K.) and grant AI175150 from the National Institutes of Health (to L.D.S.).

## Author contributions

Conceptualization, AK, LDS; Data curation, AK, WH; Formal analysis, AK, WH; Funding acquisition, LDS; Investigation, AK, WH, MK, LFJ; Methodology, AK, WH, MK, LFJ; Resources, WK, WH, MK, LFJ; Validation, AK, WH; Visualization, AK, WH; Supervision, LDS; Writing – original draft, AK, WH, LDS; Writing – review & editing, AK, WH, LDS.

## Data Sources and Availability

All materials described here are available from the corresponding author based on MTA. Sequence for the CUT&RUN analyses have been deposited to the NCBI Bioproject # PRJNA1330822.

## Declarations

The authors declare that they have no conflicts to disclose.

## MATERIALS & METHODS

### Animal studies and ethical approval

Animal studies were approved by the Institutional Animal Studies Committee (School of Medicine, Washington University in St. Louis). *Ifngr1*^−/−^ mice (003288; Jackson Laboratories), and NOD scid gamma mice (referred to as NSG) (005557; Jackson Laboratories) were bred in-house in a specific-pathogen-free animal facility on a 12:12 light-dark cycle. Male and female mice between 8 and 12 weeks of age were used for sex-matched groups used in crossing experiments. Mice were co-housed with siblings of the same sex throughout the experiments.

### Cell culture

HCT-8 cells (Human ileocecal adenocarcinoma, ATCC CCL-244) derived from a human ileocecal carcinoma were cultured in RPMI 1640 ATCC Modification medium supplemented with 10% fetal bovine serum at 37 °C in a 5% CO_2_ incubator. After trypsinization, 450,000 cells were plated on glass coverslips in 24-well plates and used for experiments. Cells were confirmed to be mycoplasma-free with the e-Myco plus Mycoplasma PCR detection kit.

#### Immunofluorescence assays & microscopy

Staining protocols have been previously described and all antibodies are listed in **Table S2** [71]. Briefly, infected HCT-8 cultures were fixed with 4% formaldehyde 8-24 hpi, permeabilized with 0.1% Triton X-100, blocked with 1% bovine serum albumin (BSA), stained with a primary antibody mixture for 1 h followed by secondary antibodies for 1 h and Hoechst for 20 min. Primary antibodies were diluted for staining: anti-HA (rat, 1:500), anti-Aldolase (ALD, rabbit, 1:1000) centrin-1 (mouse, 1:250), anti-Ty (mouse, 1:250), and anti-PH3 (rabbit, 1:500), NUP116 (rabbit, 1:100). All Alexa Fluor secondary antibodies were diluted at 1:1000 and Hoechst was diluted 1:2000. Coverslips were mounted onto microscope slides with Prolong Glass Antifade Mountant. Coverslips were visualized using a Zeiss LSM880 laser scanning confocal microscope equipped with a 63x, 1.4 N.A. Zeiss Plan Apochromat oil objective lens and Airy scan. Images were acquired with ZEN v2.1, v2.5 software and further analyzed using ImageJ. All staining experiments were performed independently 2-3 times, and at least 2-5 fields of view were imaged per coverslip.

#### Expansion Microscopy

Ultrastructure expansion microscopy (U-ExM) was performed as previously described [72]. HCT-8 cells were infected with the CENH3-3HA or CENH3-smHA parasites and fixed following the same procedures used for standard immunofluorescence assays or 16% methanol-free formaldehyde. Fixed samples were embedded overnight in a solution containing 1% acrylamide and 0.7% formaldehyde to anchor proteins and prevent crosslinking. The resulting gel was transferred into denaturation buffer and incubated at 95 °C. A second expansion gel was polymerized on ice using a monomer mixture composed of 19% sodium acrylate, 10% acrylamide, and 0.1% (1,2-Dihydroxyethylene) bisacrylamide, supplemented with 0.5% ammonium persulfate (APS) and 0.5% TEMED. Polymerized gels were denatured at 95 °C for 90 min in denaturation buffer (200 mM SDS, 200 mM NaCl, 50 mM Tris-Base, pH 9.0) and expanded overnight in ultrapure water. Expansion ratios were determined the following day by measuring gel diameters. Gels were then shrunk in PBS and stained with primary antibodies (rabbit anti-HA, rat anti-HA, or either rabbit anti-GFP antibodies at 1:200 and rat Pan-Cp, 1:500), followed by secondary antibodies (Alexa Fluor conjugates, 1:500) and Hoechst (1:500) diluted in freshly prepared 2% PBS/BSA for 3, 12, or 24 h for primary antibodies and 2 h for secondary antibodies at room temperature. After each staining step, gels were washed three times in PBS/0.1% Tween for 10 min. Stained gels were re-expanded overnight in ultrapure water before imaging. Images were acquired using a Zeiss LSM880 laser scanning confocal microscope equipped with a 63x, 1.4 N.A. Zeiss Plan-Apochromat oil objective and ZEN v2.1 or v2.5 software.

#### Immuno-electron Microscopy

Immuno-electron microscopy (EM) was performed as previously described using CENH3-3HA and CENH3-smHA parasites [73]. HCT-8 monolayers were grown in 6-well plates and reached confluence before infection with transgenic *C. parvum* oocysts. After a 20-h incubation, cultures were washed twice in DPBS and fixed for 1 h at 4 °C in freshly prepared fixative consisting of 4% paraformaldehyde and 0.05% glutaraldehyde in 100 mM PIPES supplemented with 0.5 mM MgCl₂ (pH 7.2). Fixed samples were embedded in 10% gelatin and subsequently infiltrated overnight at 4 °C with a cryoprotectant solution of 2.3 M sucrose and 20% polyvinyl pyrrolidone prepared in PIPES/MgCl₂. Blocks were trimmed, rapidly frozen in liquid nitrogen, and sectioned using a Leica Ultracut UCT7 cryo-ultramicrotome. Ultrathin sections (50 nm) were blocked for 30 min in a mixture of 5% fetal bovine serum and 5% normal goat serum, followed by incubation with rabbit anti-HA, rat anti-HA, or either of the rabbit anti-GFP antibodies (1:1000) for 1 h at room temperature. After rinsing in blocking buffer, sections were labeled for 1 h with 18 nm colloidal gold–conjugated goat anti-rabbit IgG (H+L) diluted 1:20. Labeled sections were then contrasted with 0.3% uranyl acetate in 2% methyl cellulose and imaged using a JEOL 1200 EX transmission electron microscope equipped with an AMT 8-megapixel digital camera and AMT Image Capture Engine v6.0.2. Control sections processed in parallel without primary antibody were included in all experiments.

### CRISPR/Cas9 plasmid construction

#### Plasmid construction

Primers used to generate the following plasmids were ordered from IDT and all sequences are found in **Table S3**. SnapGene (v7.1.1) and NEBuilder Assembly Tool (v2.9.0) were used to construct plasmid maps and design primers.

The CENH3-3HA tagging plasmid was generated using the previously described ABC-3HA tagging plasmid [73] by replacing the C-terminal and 3’UTR sequences with equivalent regions from the *Cp*CENH3 gene amplified from *C. parvum* cDNA to generate the tagging plasmid CENH3-3HA-Nluc-P2A-Neo^R^.

The 3HA-CENH3-GBP-3Ty tagging plasmid was created via a seven-way Gibson assembly combining a backbone generated from the TK*-*GFP-Nluc-P2A-Neo^R^-TK homology tagging plasmid previously generated in our lab [74] together with the following fragments: 1) The 5’ homology region of CENH3 (500 bp), the CENH3 CDS, a portion of the 3’ homology region of CENH3 (216 bp) were amplified from the CENH3-3HA tagging plasmid described above. 2) The 3HA-pLinker region was amplified from a TK-3HA-CENH3-Nluc-P2A-Neo^R^-TK plasmid (data not shown in this study), 3) the GBP CDS was amplified from *C. parvum* genomic DNA, 4) the pLinker-3Ty sequence was amplified from a tagging plasmid previously generated in our lab [73], and 5) the P2A skip peptide sequence was generated as a gBlock Gene Fragment from Azenta. The U6:sgTK plasmid previously generated in our lab was utilized for all experiments with TK locus replacement repair plasmids [74].

The H3.1-3HA and H3.2-3HA tagging plasmids were generated via Gibson assembly using the 3HA-CENH3-GBP-3Ty plasmid to provide a backbone, the pLinker-3HA region was amplified from the CENH3-3HA tagging plasmid described above, and the CDS regions for H3.1 and H3.2 were amplified from *C. parvum* genomic DNA.

The GBP-3HA tagging plasmid was generated using the CENH3-3HA tagging plasmid described above for the backbone and replacing the C-terminal and 3’UTR sequences with the GBP CDS sequence amplified from *C. parvum* genomic DNA.

CENH3 and GBP CRISPR/Cas9 plasmids were created via Gibson assembly of the linearized backbone amplified from the TK CRISPR/Cas9 plasmid (Addgene 122852) and an oligonucleotide with the sgRNA specific for the CENH3 or GBP gene [74]. The Eukaryotic Pathogen CRISPR guide RNA/DNA Design tool (http://grna.ctegd.uga.edu) was used to identify CENH3 and GBP sgRNA sequences.

### Amplifying transgenic parasites in mice

The transfection and amplification of parasites in immunocompromised mice were conducted as previously described [74–77]. Briefly, 1×10^8^ freshly excysted sporozoites were electroporated with 100 μg of repair plasmid and 66 μg of CRISPR/Cas9 plasmid, then resuspended in 300 μL of cold DPBS. *Ifngr1*^−/−^ mice were first orally gavaged with an 8% (wt/vol) sodium bicarbonate solution to neutralize stomach acid, followed by the administration of the electroporated sporozoites. To facilitate the selection of transgenic parasites, paromomycin (16 g/L) was provided in the drinking water 24 h post-infection. Fecal pellets were collected at intervals post-infection for luciferase assays and diagnostic PCR analysis to verify parasite growth and proper gene insertion. Oocysts were isolated from fecal material using bleach treatment, and 20,000 oocysts per mouse were employed for a subsequent round of amplification in NSG mice.

#### Luciferase assays for mouse experiments

Fecal samples collected from infected mice were analyzed with the Promega Nano-Glo luciferase assay kit, as previously described [77]. The luciferase activity was measured using the BioTek Cytation 3 cell imaging multi-mode reader. The number of relative luminescence units per milligram of feces was obtained by dividing the average values from two technical replicates by the fecal sample weight.

#### Quantifying oocyst shedding in mouse experiments

DNA was extracted from fecal pellets collected from infected mice using the QIAamp PowerFecal DNA kit, following previously established methods [74, 76, 77]. Quantitative PCR (qPCR) analysis was conducted to determine the number of oocysts at each time point based on *C. parvum* glyceraldehyde-3-phosphate dehydrogenase (GAPDH) primers and a standard curve derived from a dilution series (**S2 Table**) [74]. Genomic DNA (gDNA) amounts were quantified using the QuantStudio 3 real-time PCR system.

### Confirmation of insertions in transgenic parasites by PCR

To confirm the correct integration of the homology repair template, extracted gDNA from murine fecal samples and wild type controls were used as the template. PCR amplification was performed using 1 µL of the extracted gDNA, Q5 Hot Start high-fidelity 2X master mix or PrimeSTAR GXL premix (TaKaRa), and primers at a final concentration of 500 nM on a Veriti 96-well thermal cycler. All PCR products were resolved on a 1% agarose gel containing GelRed (1:10,000) and visualized using the ChemiDoc MP imaging system.

### Oocyst purification from fecal material and storage

The *C. parvum* isolate AUCP –1 used for transfections was purified from fecal material collected from male Holstein calves at the University of Illinois at Urbana-Champaign [78]. All procedures involving the calves received approval from the Institutional Animal Care and Use Committee (IACUC). Transgenic oocysts were purified in-house from fecal material collected from NSG mice, using a saturated sodium chloride solution for oocyst flotation [79]. Post-purification, oocysts were stored at 4°C in phosphate-buffered saline (PBS) for up to three months following fecal collection.

### Structural predictions

The protein structure of the G-quadruplex binding protein (GBP, *cgd1_*3530) was predicted using Alphafold 3 (www.alphafoldserver.com). Using ChimeraX (https://www.cgl.ucsf.edu/chimerax/), we modeled how the G-quadruplex structure formed by the *Cp*GBP repetitive binding sequence (GGTTA) [51] interacts and is stabilized by the CpGBP protein.

### Channel colocalization quantification

Co-localization analysis was performed using the Coloc2 plugin (FIJI/ImageJ), using default thresholding parameters. A 2D intensity histogram was generated as part of the Coloc2 output to visualize the pixel-wise distribution of intensities from both channels and a Pearson correlation coefficient was quantified.

### Parasite Preparation and Lysis for CUT&RUN experiments

The CENH3-3HA line of *T. gondii* [27] was obtained from the laboratory of Dr. Kami Kim at University of South Florida. *Tg*CENH3-3HA tachyzoites were propagated in human foreskin fibroblasts and harvested using established protocols [80]. *C. parvum* sporozoites (4×10^7^) and *T. gondii* tachyzoites (1×10^7^) were centrifuged at 2,500 RPM for 5 min, supernatants removed, and parasites were resuspended in 2 mL of cold parasite lysis buffer (10 mM HEPES pH 7.9, 1.5 mM MgCl2, 10 mM KCl, 0.65% NP40 (supplemented with IGEPAL) and 0.5 mM PMSF (supplemented with cOmplete EDTA free protease inhibitor cocktail and sterile distilled H_2_O), incubated for 10 min on ice, and centrifuged at 3,000 RPM for 10 min at 4°C (previously described in [81, 82]).

### CUT&RUN Processing, Library Preparation, Quality Control & Sequencing

Following lysis treatment, the lysis buffer was removed, and the nuclei pellet was washed following the CUT&RUN Kit Version 4 protocol (EpiCypher, Inc, Catalog No. 14-1048). Isolated nuclei were incubated with either 1.0 μg of rat anti-HA antibody, 0.5 μg of rabbit anti-H3K4me3 positive control antibody, or 0.5 μg of mouse anti-IgG negative control antibody. Chromatin digestion and release steps were performed according to the EpiCypher protocol, and 5 ng of CUT&RUN enriched DNA was used for library preparation according to the CUT&RUN Library Prep Kit User Manual Version 1.5 (EpiCypher, Inc, Catalog No. 14-1001). The concentrations of CUT&RUN library samples were determined via Qubit and quality control were assessed with an Agilent 4200 TapeStation.

### CUT&RUN Analysis & Visualization

Multiple CUT&RUN runs were preformed to obtain biological replicates of each sample. DNA products were sequenced on an Illumina NovaSeq 6000 using 150-bp paired-ends at the GTAC sequencing core at Washington University in St. Louis. Sequence reads obtained from Illumina were trimmed for adapter sequences and poor sequence quality (phred score < 25 and trimmed length < 50) using fastp v0.23.4. The trimmed reads were mapped to the reference *C. parvum* IOWA-CDC genome using BWA-MEM v0.7.17 [83]. Duplicate reads were marked and removed in the bam files using the MarkDuplicates algorithm of picard v3.1.1 (https://broadinstitute.github.io/picard/) with REMOVE_DUPLICATES=true, CREATE_INDEX=true, and ASSUME_SORTED=true. Reads with mapping lengths less than 50 bp were further removed from the bam files using SAMtools v1.7 (http://samtools.sourceforge.net/). Genome coverage and sequence depth were estimated using the mpileup algorithm of SAMtools. The read depth of each locus was normalized by dividing by the total number of counts for all loci, then scaled by multiplying by the number of loci in the genome. Input counts were subtracted from negative control counts. Data was visualized by a R package ‘ggplot2’ (https://www.rdocumentation.org/packages/ggplot2). Motifs within centromeric windows were identified using MEME v5.5.8 (https://meme-suite.org/meme/tools/meme). Chromosomal distributions of intergenic regions, centromeric windows, and motifs were visualized with Circos plot generated in TBtools-II v2.311 (https://github.com/CJ-Chen/TBtools-II).

## Supporting Information

**S1 Fig.** Tagged versions of the H3.1, H3.2, and CENH3 and GBP genes were inserted into the tk locus in *Cryptosporidium parvum*.

**S2 Fig.** The GBP and CENH3 expression overlaps in *Cryptosporidium parvum*.

**S3 Fig.** The GBP and CENH3 gene were successfully tagged with HA tag and/or a spaghetti monster HA tag in C*ryptosporidium parvum*.

**S4 Fig.** Additional images of centrin and tubulin staining in *Cryptosporidium parvum*.

**S5 Fig.** Genome-wide distribution of CENH3 in *Toxoplasma gondii*.

**S6 Fig.** Genome-wide distribution of CENH3 in *Cryptosporidium parvum*.

**S1 Table.** Transgenic *Cryptosporidium* strains used in this study.

**S2 Table.** Key Resources used in the experiments.

**S3 Table.** Oligonucleotides and plasmids used in this study.

**S1 Dataset** Information of sequencing samples from the Cut&Run and summary statistics of the data.

**S2 Dataset** Genomic location of the CENH3 sites within coding regions.

**S3 Dataset** Genomic location of the CENH3 sites within intergenic regions.

