## Supporting information for "Evidence for holocentric centromeres in the early branching apicomplexan parasite *Cryptosporidium parvum*"

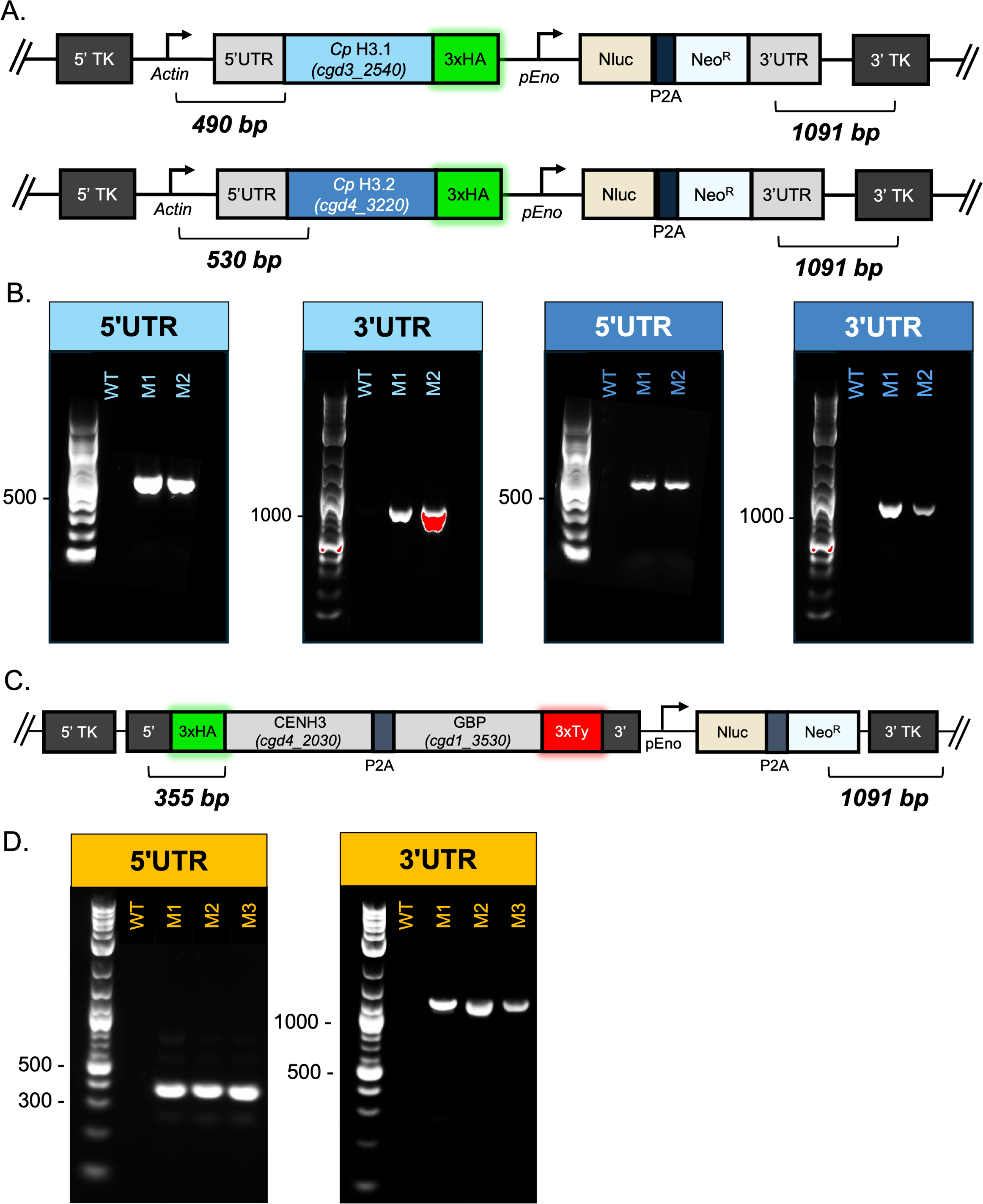

**S1 Fig. Tagged versions of the H3.1, H3.2, and CENH3 and GBP genes were inserted into the tk locus in *Cryptosporidium parvum*.** (A) A diagram of the TK-H3.1-3HA-Nluc-Neo-TK and TK-H3.2-3HA-Nluc-Neo-TKtargeting vector and the expected sizes for the 5’ insertion site (490 and 530 bp, respectively) and 3’ insertion site (1091 bp) amplicons used to confirm proper insertion by PCR. (B) PCR analysis of H3.1-3HA and H3.2-3HA oocysts amplified in GKO mice. DNA purified from fecal samples collected from each mouse and a wild type (WT) control was used to amplify products specific for the 5′ and 3’ insertion sites of the integrated construct. (C) A diagram of the TK-3HA-CENH3-P2A-GBP-3Ty-Nluc-Neo-TK targeting vector and the expected sizes for the 5’ UTR (355 bp) and 3’ UTR (1091 bp) amplicons used to confirm proper insertion by PCR. (D) PCR analysis of TK-3HA-CENH3-P2A-GBP-3Ty-TK oocysts amplified in three NSG mice. DNA purified from fecal samples collected from each mouse (M1-3) and a wild type (WT) control was used to amplify products specific for the 5′ and 3’ insertion sites of the integrated construct.

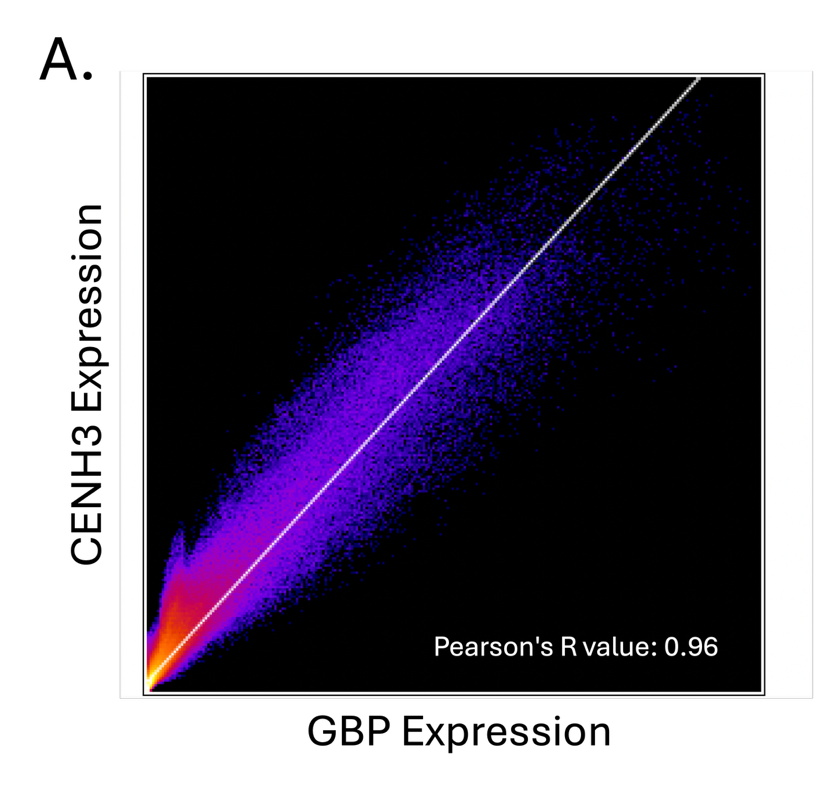

**S2 Fig. Quantification of the overlap in expression of GBP and CENH3 signals in *Cryptosporidium parvum.*** (A) The microscopy image shown in Fig. 3F was used to quantify the overlap between the 488‑nm channel (detecting CENH3‑3HA with an anti‑HA primary antibody and an Alexa Fluor 488 secondary antibody) and the 568‑nm channel (detecting GBP‑3Ty via an anti‑Ty primary antibody and an Alexa Fluor 568 secondary antibody). A 2D intensity histogram generated using the Coloc2 plugin in FIJI/ImageJ displays the pixel‑wise relationship between both channels. The accompanying text indicates the classical Pearson correlation coefficient calculated from the pixel‑intensity correlation, reflecting the degree of co‑localization between CENH3 and GBP signals.

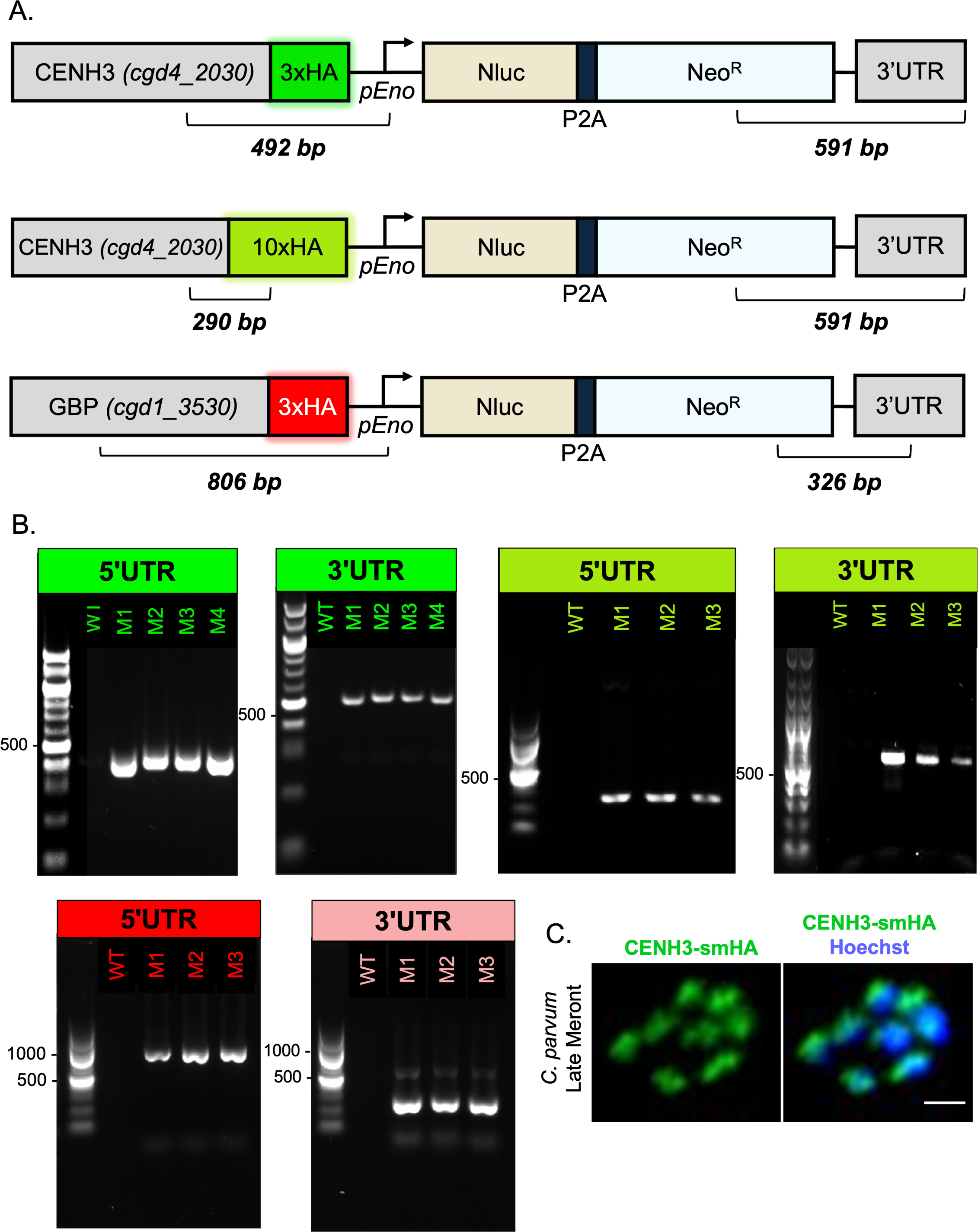

**S3 Fig. The GBP and CENH3 gene were successfully tagged with HA tag and/or a spaghetti monster HA tag in *Cryptosporidium parvum.*** (A) A diagram of the CENH3-3HA-Nluc-Neo, CENH3-smHA-Nluc-Neo, and GBP-3HA-Nluc-Neo targeting vectors and the expected sizes for the 5’insertion site (492, 290, and 806 bp, respectively) and 3’insertion site (591 and 326 bp, respectively) amplicons used to confirm proper insertion by PCR. (B) PCR analysis of CENH3-3HA, CENH3-smHA and GBP-3HA oocysts amplified in NSG mice. DNA purified from fecal samples collected from each mouse and a wild type (WT) control was used to amplify products specific for the 5′ and 3’ insertion sites of the integrated construct. (C) HCT-8 cells were infected with CENH3-smHA *Cp* parasites, fixed at 22 hpi, and stained with rat anti-HA and VVL-Biotin, followed by secondary antibodies Alexa Fluor 488 goat anti-rat IgG and Alexa Fluor 647 Streptavidin, followed by Hoechst staining. Scale bars, 1 μm.

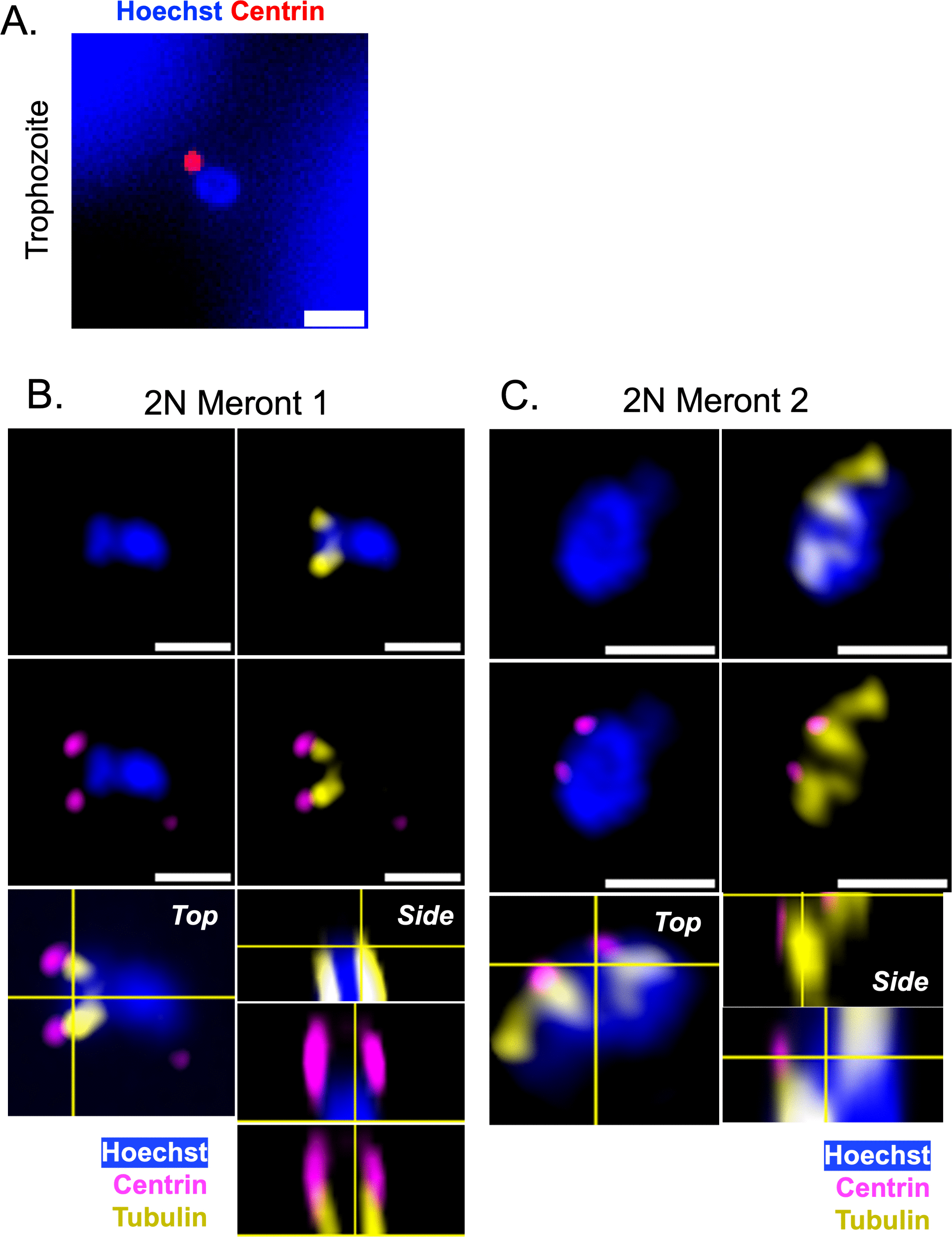

**S4 Fig. Additional images of centrin and tubulin staining in *Cryptosporidium parvum.*** (A) HCT-8 cells were infected with excysted wildtype *C. parvum* sporozoites for 2 h, then washed twice to remove extracellular parasites. Cultures were fixed at 30-min intervals between 6-9 hpi to identify trophozoites before the first mitotic division (nucleus with one centrin) by widefield microscopy. Cultures were stained with rabbit anti-centrin-1, the secondary antibody Alexa Fluor 568 goat anti-rabbit IgG, and lastly Hoechst. Scale bars, 1 μm. (B, C) HCT-8 cells were infected with wildtype parasites, fixed at 6.5 hpi to capture early meronts (2 nuclei), and stained with rabbit anti-centrin-1 and mouse anti-tubulin, followed by the secondary antibodies Alexa Fluor 488 goat anti-mouse IgG, Alexa Fluor 568 goat anti-rabbit IgG, and Hoechst. Meronts shown in panels B and C are identical to the meronts found in Fig 4C and D, respectively, but additional single-channel panels are shown here. Images were acquired as Z-stacks using LSCM-A and are presented with orthogonal views. Scale bars, 1 μm.

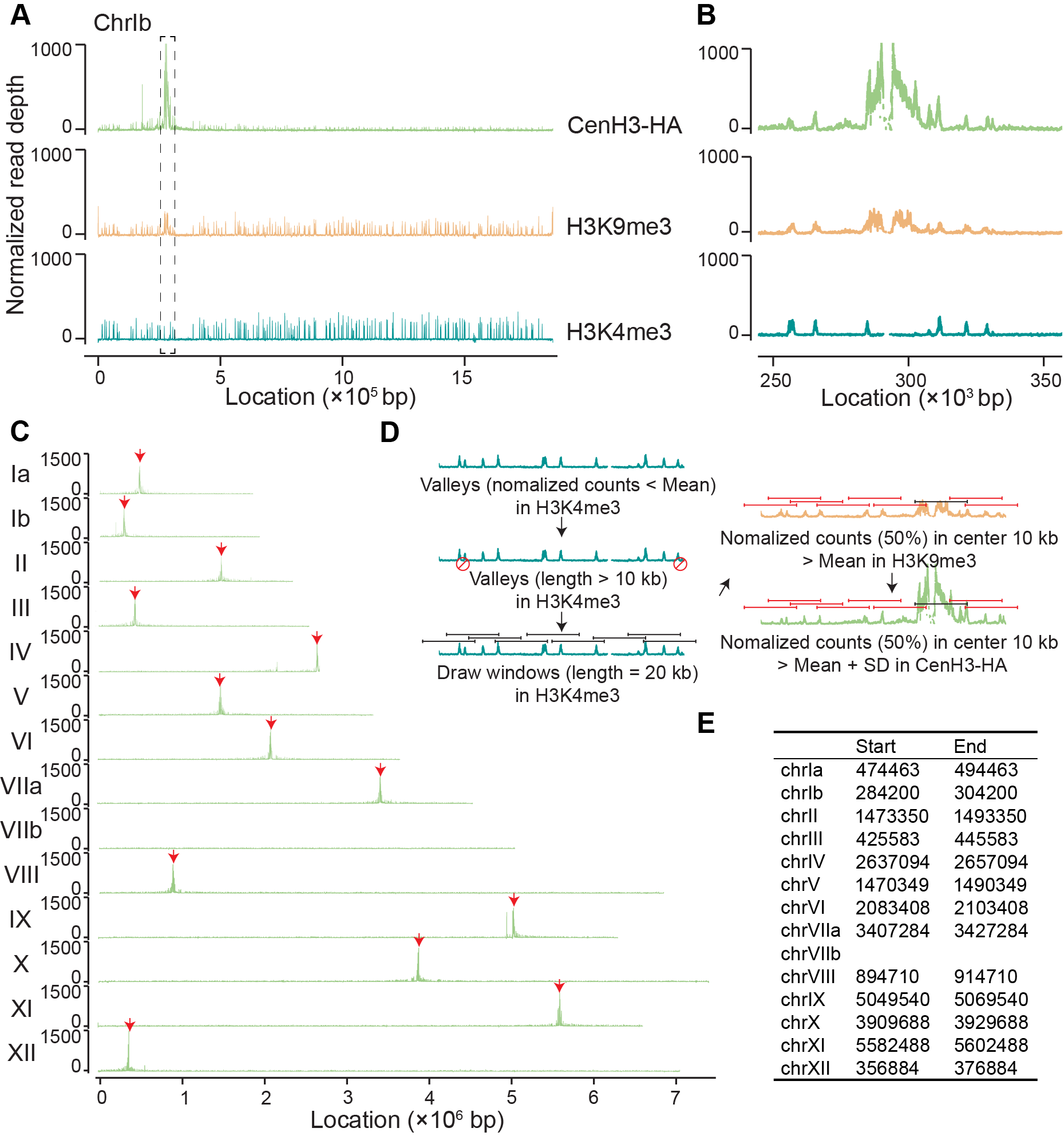

**S5 Fig. Genome-wide distribution of CENH3 in *Toxoplasma gondii****.* (A) Genome browser view of *T. gondii* chromosome Ib showing CUT&RUN signals for CENH3-HA (HA antibody), H3K9me3, and H3K4me3. CENH3 signal enrichment positively correlates with H3K9me3 and inversely with H3K4me3. Peaks of CENH3 enrichment are indicated (dashed box). (B) Zoom in genome browser view of the dashed box in (A). (C) Genome browser view of all 13 *T. gondii* chromosomes showing CUT&RUN signals for CENH3-HA, with CENH3 peaks indicated (red arrows). (D) Schematic workflow for identifying centromeric windows. (E) List of centromeric windows identified in (D), corresponding to the peaks observed in the genome browser.

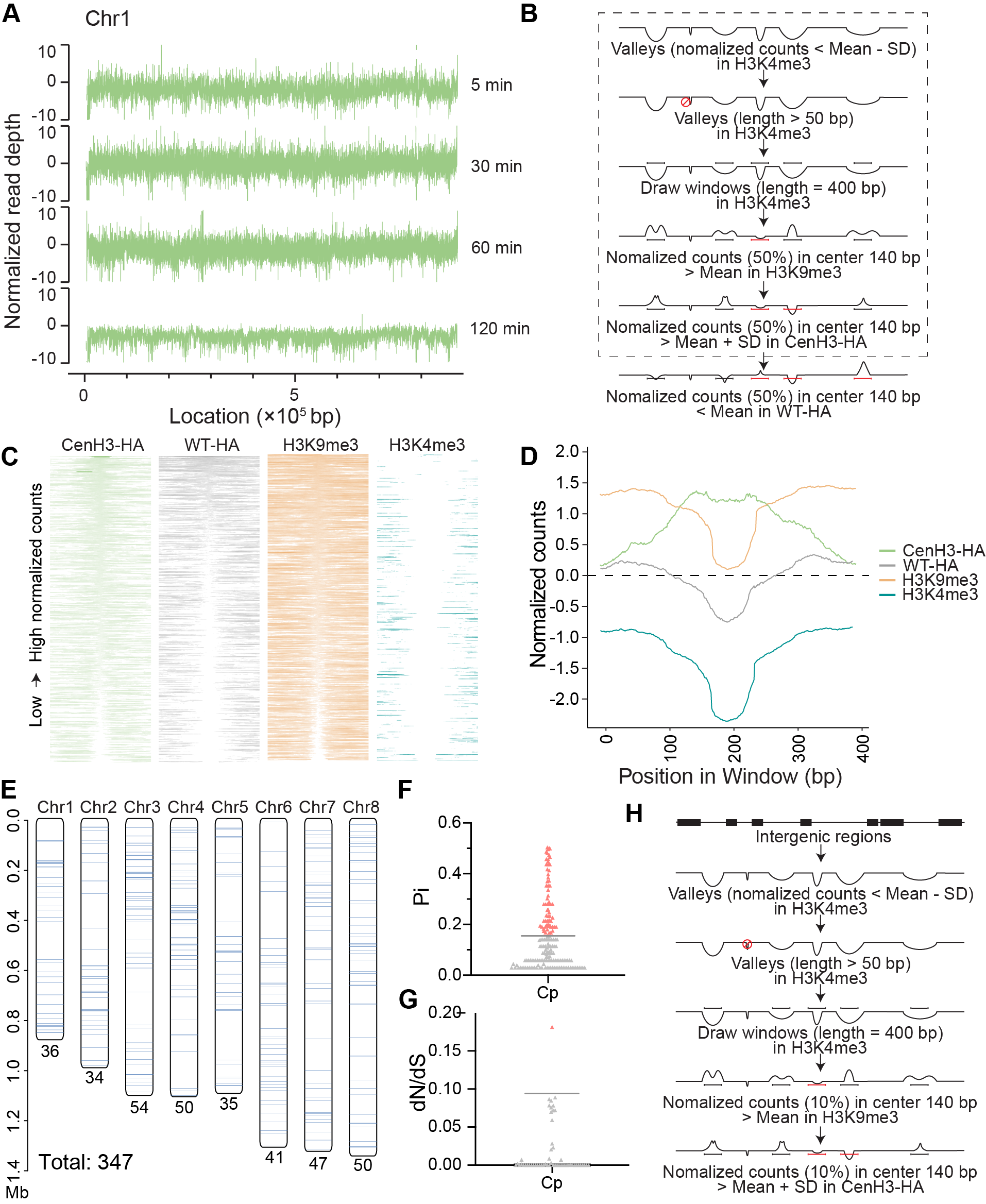

**S6 Fig. Genome-wide distribution of CENH3 in *Cryptosporidium parvum.***

(A) Genome browser view of chromosome 1 showing CENH3-HA CUT&RUN profiles at different digestion times (5, 30, 60, and 120 min). (B) Schematic workflow for identifying centromeric windows (dashed box), including additional filtering based on WT-HA signal patterns. (C) Heatmaps of CUT&RUN signals within a 400 bp window around all 423 CENH3-enriched windows identified using the stepwise strategy showing in (B, dashed box). CUT&RUN signals for CENH3-HA *C. parvum* captured with HA antibody, wild type *C. parvum* captured with HA antibody (referred to as WT-HA), and wild type C. parvum captured with H3K9me3, and H3K4me3 antibodies. Each row represents an individual centromeric site, ranked by signal intensity. (D) Average signal profiles for CENH3-HA (green), WT-HA (gray), H3K9me3 (orange), and H3K4me3 (teal) across the 400 bp window around the 423 CENH3-enriched windows. (E) Chromosomal distribution of the 347 CENH3-enriched windows of *C. parvum* chromosomes (Chr#) with the additional filtering based on removing WT-HA signal. Numbers below each chromosome indicate the number of windows. (F, G) Evolutionary constraint analysis of CENH3-associated regions using nucleotide diversity (Pi) and dN/dS values. Comparisons were performed across *C. parvum* IIa and IId isolates (referred to as Cp). (F) Horizontal lines indicate the mean Pi values of genome-wide 400-bp sliding windows, used as a reference for comparison. Approximately 82% of CENH3-associated regions showed reduced Pi values when compared to the genome as a whole. (G) Horizontal lines indicate the mean dN/dS value of all orthologous genes with *P* < 0.05, used as a reference for comparison. Only one gene with in the CENH3-associated regions showed elevated dN/dS relative to the genome-wide distribution. (H) Schematic workflow for identifying centromeric windows located in intergenic regions.

**S1 Table.** Transgenic *Cryptosporidium* strains used in this study.

| **Line** | **Genotype** | **Use** | **Endogenous or second copy** | **Source** |
| --- | --- | --- | --- | --- |
| H3.1-3HA | TK-H3.1-3HA-Nluc-P2A-Neo^R^-TK | Epitope tagging for localization | Second copy | This study |
| H3.2-3HA | TK-H3.2-3HA-Nluc-P2A-Neo^R^-TK | Epitope tagging for localization | Second copy | This study |
| CENH3-3HA | CENH3-3HA-Nluc-P2A-Neo^R^ | Epitope tagging for localization, ExM, EM | Endogenous | This study |
| GBP-3HA | GBP-3HA-Nluc-P2A-Neo^R^ | Epitope tagging for localization | Endogenous | This study |
| 3HA-CENH3-GBP-Ty | TK-3HA-CENH3-P2A-GBP-3Ty-Nluc-P2A-Neo^R^-TK | Epitope tagging for localization | Second copy | This study |
| CENH3-smHA | CENH3-smHA-Nluc-P2A-Neo^R^ | CUT&RUN, ExM, EM | Endogenous | This study |

**S2 Table.** Key Resources used in the experiments.

| **REAGENT or RESOURCE** | **SOURCE** | **IDENTIFIER** |
| --- | --- | --- |
| **Antibodies** | | |
| Rat monoclonal anti-HA (3F10) (IFA, ExM, and EM) | Sigma-Aldrich | Cat# 11867423001 |
| Rabbit polyclonal anti-HA (SG77) (ExM and EM only) | Novus Biologicals | Cat# 71-5500 |
| Polyclonal Rabbit anti-GFP (ExM and EM only) | Origene | TP401 |
| Polyclonal Rabbit anti-GFP (ExM and EM only) | ThermoFisher | Cat# A-11122 |
| Rabbit anti-phosphorylated Histone H3 (PH3) | Sigma-Aldrich | Cat# H0412 |
| Rabbit anti-Centrin | Kerafast | Cat# EBC004 |
| Mouse anti-tubulin (12G10) | Developmental Studies Hybridoma Bank | Antibody ID# AB_1157911 |
| Rabbit *Plasmodium falciparum* nuclear pore (NUP116) | The Laboratory of Dr. Artur Scherf [1] | *Pf*Nup116 |
| Mouse anti-Ty | In house hybridoma | mAb Clone BB2 |
| *Vicia Villosa* Lectin (VVL), Biotinylated | Vector Labs | Cat# B-1235-2 |
| Rat PanCp (ExM only) | In house antisera [2] |  |
| Alexa Fluor 488 goat anti-rat IgG (H+L) | Thermo Fisher Scientific | Cat# A11006 |
| Alexa Fluor 488 goat anti-rabbit IgG (H+L) (IFA and ExM) | Thermo Fisher Scientific | Cat# A11034 |
| Alexa Fluor 568 goat anti-rat IgG (H+L) (IFA and ExM) | Thermo Fisher Scientific | Cat# A11077 |
| Alexa Fluor 568 goat anti-rabbit IgG (H+L) | Thermo Fisher Scientific | Cat# A11011 |
| Alexa Fluor 647 goat anti-mouse IgG (H+L) | Thermo Fisher Scientific | Cat# A21235 |
| Alexa Fluor 647 Streptavidin | Thermo Fisher Scientific | Cat# S21374 |
| 18 nm colloidal gold–conjugated goat anti‑rabbit IgG (H+L) (EM only) | Jackson ImmunoResearch Laboratories Inc. | Cat# 111‑215‑144 |
| Hoechst | Thermo Fischer Scientific | Cat# C10637 |
| H3K9me2 | Sigma-Aldrich | Cat# SAB4200609 |
| H3K9me3 | Abcam | Cat# ab8898 |
| HA Tag CUTANA™ CUT&RUN Antibody | Epicypher | Cat# 13-2010 |
| ***Cryptosporidium* & Bacterial Strains** | | |
| *Cryptosporidium parvum* AUCP-1 isolate | Laboratory of William Witola | N/A |
| NEB 5-alpha Competent E. coli (High Efficiency) | New England Biolabs | Cat# C2987H |
| **Chemicals, Reagents & Supplies** | | |
| Sodium taurocholate hydrate | Sigma-Aldrich | Cat# 86339 |
| Sodium chloride | Sigma-Aldrich | Cat# S7653-250G |
| Tween-20 | Sigma-Aldrich | Cat# 11332465001 |
| Fetal bovine serum | Gibco | Cat# 10-082-147 |
| Paromomycin sulfate salt | Sigma-Aldrich | Cat# P9297 |
| Bovine serum albumin | Sigma-Aldrich | Cat# A7030 |
| Poly-L-Lysine solution (0.01%) | Sigma-Aldrich | Cat# P4707 |
| Triton X-100 | Thermo Fisher Scientific | Cat# BP151 |
| ProLong Glass Antifade Mountant | Thermo Fisher Scientific | Cat# P36984 |
| Formaldehyde (methanol-free), Ultrapure EM Grade | Polysciences, Inc. | Cat# 04018-1 |
| GelRed nucleic acid gel stain | Biotium | Cat# 41003-1 |
| Sodium bicarbonate | ATCC | Cat# 30-2002 |
| Carbenicillin disodium salt | Sigma-Aldrich | Cat# C3416 |
| Fetal bovine serum (FBS) | Sigma-Aldrich | Cat# F6178 |
| RPMI 1640 ATCC Modification medium | Thermo Fisher Scientific | Cat# A14091-01 |
| Dulbecco’s Phosphate-Buffered Saline (DPBS) | Invitrogen | Cat# 15575020 |
| Falcon 15 mL/50 mL Polystyrene Centrifuge Tubes | Corning Costar | Cat# 352099, 430290 |
| HEPES-KOH | Sigma-Aldrich | Cat# H0527 |
| MgCl2 | Sigma-Aldrich | Cat# 208337 |
| KCl | Sigma-Aldrich | Cat# P3911 |
| IGEPAL CA-630 | Sigma-Aldrich | Cat# I8896 |
| cOmplete EDTA-free Protease Inhibitor Cocktail | Sigma-Aldrich | Cat# 11873580001 |
| **Commercial Assays** | | |
| e-Myco plus Mycoplasma PCR detection kit | Boca Scientific | Cat#: 25237 |
| QIAamp DNA Mini kit | QIAGEN | Cat# 51306 |
| QIAamp Powerfecal Pro DNA kit | QIAGEN | Cat# 51804 |
| TB Green Advantage qPCR premix | Takara Bio | Cat# 639676 |
| Gibson Assembly Cloning kit | New England Biosciences | Cat# E5510S |
| Q5 Site-directed Mutagenesis kit | New England Biosciences | Cat# E0554S |
| Q5 Hot Start High-Fidelity 2X master mix | New England Biosciences | Cat# M0494S |
| SF Cell Line 4D-Nucleofector X Kit L | Lonza | Cat# V4XC-2024 |
| Nano-Glo Luciferase Assay kit | Promega | Cat# N1120 |
| CUTANA™ ChIC/CUT&RUN Kit | Epicypher | Cat# 14-1048 |
| CUTANA™ CUT&RUN Library Prep Kit | Epicypher | Cat# 14-1001 |
| **Experimental Models: Cell Lines** | | |
| Human: HCT-8 | ATCC | CCL-244 |
| **Experimental Models: Organisms/Strains** | | |
| Mouse: Ifngr1-/- (C57BL/6 background) | Jackson Laboratories | Cat# 003288 |
| Mouse: Nod scid gamma (NSG) | Jackson Laboratories | Cat# 005557 |
| **Oligonucleotides** | | |
| Primer: *C. parvum* GAPDH forward: CGGATGGCCATACCTGTGAG | [2] | N/A |
| Primer: *C. parvum* GAPDH reverse: GAAGATGCGCTGGGAACAAC | [2] | N/A |
| **Recombinant DNA** | | |
| Plasmid: TK-H3.1-3HA -Nluc-P2A-neo-TK | This paper | N/A |
| Plasmid: TK-H3.2-3HA -Nluc-P2A-neo-TK | This paper | N/A |
| Plasmid: CENH3-3HA-Nluc-P2A-neo | This paper | N/A |
| Plasmid: GBP-3HA-Nluc-P2A-neo | This paper | N/A |
| Plasmid: TK-3HA-CENH3-P2A-GBP-3Ty-Nluc-P2A-neo-TK | This paper | N/A |
| Plasmid: CENH3-smHA-Nluc-P2A-neo | This paper | N/A |
| Plasmid: pACT1:Cas9, U6:sgTK | [2] | Cat# 122852 |
| Plasmid: pACT1:Cas9, U6:sgCENH3 | This paper | N/A |
| Plasmid: pACT1:Cas9, U6:sgGBP | This paper | N/A |
| **Instruments, Software, and Algorithms** | | |
| GraphPad Prism 10 | GraphPad Software | <https://www.graphpad.com/> |
| QuantStudio Design & Analysis Software | Thermo Fisher Scientific | <https://www.thermofisher.com/us/en/home/global/forms/life-science/quantstudio-3-5-software.html> |
| FIJI (ImageJ) |  | <https://fiji.sc/> |
| Zeiss Observer Z1 inverted microscope with a Colibri LED illumination for multi-color epifluorescence | Zeiss | <https://www.zeiss.com/microscopy> |
| Hamamatsu ORCA-ER CCD Camera | Hamamatsu | <https://www.hamamatsu.com/us/en/product/cameras> |
| ZEN 2.5 software | Zeis | <https://www.zeiss.com/microscopy/en/products/software/zeiss-zen> |
| Zeiss Axioskop Mot Plus fluorescence microscope | Zeiss | <https://www.zeiss.com/microscopy> |
| AxioCam MRm monochrome digital camera and Axiovision software | Zeiss | <https://www.zeiss.com/microscopy/en/products/cameras> |
| Cytation 3 cell imaging multi-mode reader | Biotek | <https://www.agilent.com/en/product/microplate-instrumentation/microplate-readers/multimode-microplate-readers/biotek-cytation-hybrid-multimode-reader-1623197> |
| Veriti 96-well thermal cycler | Applied Biosystems | <https://www.thermofisher.com/order/catalog/product/4375305> |
| Qubit 4 Fluorometer | Invitrogen | <https://www.thermofisher.com/order/catalog/product/Q33226> |
| Alphafold 3 | Alphafold server | https://alphafoldserver.com/ |
| ChimeraX | UCSF ChimeraX | <https://www.cgl.ucsf.edu/chimerax/> |
| fastp v0.23.4 | Github | <https://github.com/OpenGene/fastp> |
| BWA-MEM v0.7.17 | BWA | <https://github.com/lh3/bwa> |
| picard v3.1.1 | Github | <https://broadinstitute.github.io/picard/> |
| SAMtools v1.7 | SAMtools | <http://samtools.sourceforge.net/> |
| ggplot2 | R | <https://www.rdocumentation.org/packages/ggplot2> |
| MEME v5.5.8 | MEME suite | <https://meme-suite.org/meme/tools/meme> |
| TBtools-II v2.311 | Github | <https://github.com/CJ-Chen/TBtools-II> |

**S3 Table.** Oligonucleotides and plasmids used in this study.

| **Usage** | **Full Plasmid name** | **Oligo name** | **Sequence (5’ – 3’)** | **Purpose** | **Source** |
| --- | --- | --- | --- | --- | --- |
| Inserting an epitope tagged cgd3_2540 (H3.1) & cgd4_3220 (H3.2) into the TK locus | TK- cgd3_2540 (H3.1) -3HA-Nluc-P2A-neo-TK & TK- cgd4_3220 (H3.2)-3HA-Nluc-P2A-neo-TK | TK-cgd3_2540 BB Gib F | agattatgcctaatgttttttttaccaaataaatagctc | Amplify backbone from TK-GFP- Nluc-P2A-neo-TK plasmid for building TK- cgd3_2540 (H3.1) -3HA-Nluc-P2A-neo-TK by Gibson assembly | IDT |
|  |  | TK-cgd3_2540 BB Gib R | ttagaaacaaggtatgagtgatattattgctttaaag |  |  |
|  |  | TK-cgd4_3220 BB Gib F | agattatgccTAATGTTTTTTTTACCAAATAAATAGCTC | Amplify backbone from TK-GFP- Nluc-P2A-neo-TK plasmid for building TK- *cgd4_3220 (H3.2)-*3HA-Nluc-P2A-neo-TK by Gibson assembly |  |
|  |  | TK-cgd4_3220 BB Gib R | ttcaagcattggtatgagtgatattattgctttaaag |  |  |
|  |  | cgd3_2540-3xHA Gib F | gggtgaaagggctagcaagggctcgggc | Amplify pLinker-3xHA region from the CENH3-3HA-Nluc-P2A-neo plasmid for building TK- *cgd4_3220 (H3.2)-*3HA-Nluc-P2A-neo-TK by Gibson assembly |  |
|  |  | cgd3_2540-3xHA Gib R | AAAAAAAACATTAggcataatctggaacatcgtaaggatacg |  |  |
|  |  | cgd4_3220-3xHA Gib F | tgagaggtgagctagcaagggctcgggc | Amplify pLinker-3xHA region from the CENH3-3HA-Nluc-P2A-neo plasmid for building TK- *cgd4_3220 (H3.2)-*3HA-Nluc-P2A-neo-TK by Gibson assembly |  |
|  |  | cgd4_3220-3xHA Gib R | AAAAACATTAggcataatctggaacatcgtaaggatacg |  |  |
|  |  | TK-cgd3_2540 Gib F | cactcataccttgtttctaattagttcttcagatc | Amplify cgd3_2540 (H3.1) gene from *Cryptosporidium* DNA for building TK- *cgd3_2540 (H3.1)-*3HA-Nluc-P2A-neo-TK by Gibson assembly |  |
|  |  | TK-cgd3_2540 Gib R | ccttgctagccctttcacccctaattcttc |  |  |
|  |  | TK-cgd4_3220 Gib F | cactcataccaatgcttgaaagtacaagtgtgg | Amplify cgd4_3220 (H3.2) gene from *Cryptosporidium* DNA for building TK- *cgd4_3220 (H3.2)-*3HA-Nluc-P2A-neo-TK by Gibson assembly |  |
|  |  | TK-cgd4_3220 Gib R | ccttgctagctcacctctcaccacggatac |  |  |
|  |  | Cp-TK F2 | ACCATCTAACACAAAGAATTCAACA | To check correct gene insertion from DNA purified from mouse feces. | [2]; IDT |
|  |  | 3HA R | ggcataatctggaacatcgtaagg |  |  |
|  |  | Neo F1 | GCTGAAGAACTTGGTGGTGA |  |  |
|  |  | Cp TK-R | CCTACCATAAGCATCACCAGT |  |  |
| Epitope tagging of cgd4_2030 (CENH3) | CENH3 (cgd4_2030)-3HA-Nluc-P2A-neo | cgd4_2030 CENH3 BB Gib F | ctcggtacccGCTAGCAAGGGCTCGGGC | Amplify backbone from ABC-3HA-Nluc-P2A-neo plasmid for building CENH3 (cgd4_2030)-3HA-Nluc-P2A-neo by Gibson assembly | IDT |
|  |  | cgd4_2030 CENH3 BB Gib R | ccttgctagcGGGTACCGAGCTCGAATTCAC |  |  |
|  |  | cgd4_2030 CENH3 5' homology insert Gib F | ctcggtacccAACGAATAATAGAGCAATTAGTAG | Amplify cgd4_2030 (CENH3) 5' homology and 3'UTR from *Cryptosporidium c*DNA for building CENH3 (cgd4_2030)-3HA-Nluc-P2A-neo by Gibson assembly |  |
|  |  | cgd4_2030 CENH3 5' homology insert Gib R | ccttgctagcAATAAAGTCTCCATACCTGC |  |  |
|  |  | cgd4_2030 CENH3 3'UTR insert Gib F | atctgaacaaacggaggagtcttgttttttttaccaaataaatagctcc |  |  |
|  |  | cgd4_2030 CENH3 3'UTR insert Gib R | CCAACTCGGCCATCCACGCGccgctactaccgaccgttta |  |  |
|  | CENH3 Cas9 generation and PAM mutation | CENH3 gRNA2 | gTACAGGTATATAGTACTCCTgttttagagctagaaatagcaagt | Generate Cas9 backbone using DNA from the pCRISPR-Enolase-gRNA plasmid | IDT |
|  |  | U6 F | CCCAACACTTAACCTTTCAGT |  |  |
|  |  | CENH3 PAM MUT gRNA2 F1 | agtactcctagttcaatttaattca | Mutate the PAM sequence in the CENH3 tagging plasmid so that it is not recognized by the Cas9 |  |
|  |  | CENH3 PAM MUT gRNA2 R1 | atatacctgtaatatatgttaattttagtt |  |  |
|  | Gene insertion confirmation | CenH3 5'UTR check F2 | AGTAGCCCCATTAGCCCAAG | To check correct gene insertion from DNA purified from mouse feces. | [2]; IDT |
|  |  | 3HA-R | ggcataatctggaacatcgtaagg |  |  |
|  |  | CenH3 3'UTR check R | GGCGGGGATAATTCATGTTT |  |  |
|  |  | NeoF1 | GCTGAAGAACTTGGTGGTGA |  |  |
| Epitope tagging of cgd1_3530 (GBP1) | GBP (cgd1_3530)-3HA-Nluc-P2A-neo | LacZ F2 | GGGGATCCTCTAGAGTCGAC | Generate AmpR region of the plasmid backbone from the Enolase-3HA-Nluc-neo-gRNA2 plasmid | [3]; IDT |
|  |  | LacZ R1 | GGGTACCGAGCTCGAATT |  |  |
|  |  | Linker F1 | gctagcaagggctcgggc | Generate the selection casette region of the plasmid backbone from the Enolase-3HA-Nluc-neo-gRNA2 plasmid |  |
|  |  | Neo R2 | caattgtcagaagaattcgt |  |  |
|  |  | Gbp 5' homo Gib F1 | ctcggtacccTGAATTAAATAATACTACATTACTTGATC | Amplify cgd1_3530 (GBP1) 5' homology and 3'UTR from *Cryptosporidium* gDNA for building GBP1 (cgd1_3530)-3HA-Nluc-P2A-neo by Gibson assembly | IDT |
|  |  | Gbp 5' homo Gib R1 | gagcccgagcccttgctagcCTGGTCGATTCTAACCAATA |  |  |
|  |  | Gbp 3' UTR Gib F1 | ctgacaattgATTAATTCAAAATTACTTTAAAGTCAATTTTTTG |  |  |
|  |  | Gbp 3' UTR Gib R1 | cttgctagcAATTGAACTTTCTATTTTTTTTCATTG |  |  |
|  | GBP Cas9 generation and PAM mutation | Tracer RNA F1 | gttttagagctagaaatagcaag | Generate Cas9 backbone using DNA from the pCRISPR-Enolase-gRNA plasmid | [3]; IDT |
|  |  | U6 F | CCCAACACTTAACCTTTCAGT |  |  |
|  |  | cgd1_3530_gRNA1_105 Linker | ctgaaaggttaagtgttgggGAGGCTATGGAATGAGAACAgttttagagctagaaatagc | gRNA linker sequence for Gibson assembly with Cas9 backbone | IDT |
|  |  | GBP PAM MUT gRNA1 F1 | ATGAGAACAAAGACTCATGCT | Mutate the PAM sequence in the GBP1 tagging plasmid so that it is not recognized by the Cas9 |  |
|  |  | GBP PAM MUT gRNA1 R1 | TCCATAGCCTCTGTTATTCCA |  |  |
|  | Gene insertion confirmation | Gbp Insert F1 | GAGCTATCAATGAATTAAATAATAC | To check correct gene insertion from DNA purified from mouse feces. | [2]; IDT |
|  |  | pEno R | tagagtttggtgtgcaggcg |  |  |
|  |  | NeoF1 | GCTGAAGAACTTGGTGGTGA |  |  |
|  |  | Gbp Insert R2 | GAGTGGTGTTTTACCATGGAATTGATATCATTATC |  |  |
| Inserting an epitope tagged cgd4_2030 (CENH3) & cgd1_3530 (GBP) into the TK locus | Generating TK-3HA-CENH3-P2A-GBP-3TY-TK tagging plasmmid | CenH3-Gbp Backbone F | GGTCGGTAGTAGCGGaggtggggaaactaaatata | Amplify backbone from TK-3HA-CENH3-Nluc-P2A-neo-TK plasmid (not featured in MS) for building TK-3HA-CENH3-P2A-GBP-3TY-TK by Gibson assembly | IDT |
|  |  | CenH3-Gbp Backbone R | ACAACCAAAATATTCggtatgagtgatattattgc |  |  |
|  |  | CenH3 5'UTR-3xHA-pLinker F | aatatcactcataccGAATATTTTGGTTGTTTAATTAAGATATTATTCATGCATGAAACG | Amplify 3HA-pLinker region from TK-3HA-CENH3-Nluc-P2A-neo-TK plasmid (not featured in MS) for gibson assembly |  |
|  |  | CenH3 5'UTR-3xHA-pLinker R | TTTAGAACGAGCCATcagctgggtcgagcccga |  |  |
|  |  | CenH3 CDS F | ggctcgacccagctgATGGCTCGTTCTAAAACTTC | Amplify CENH3 gene from *Cryptosporidium* cDNA for gibson assembly |  |
|  |  | CenH3 CDS R | agtagctccgcttccAATAAAGTCTCCATACCTGC |  |  |
|  |  | Gbp CDS F | gagaaccctggacctATGGTATCTGATAAAAACTGC | Amplify GBP gene from *Cryptosporidium* gDNA for gibson assembly |  |
|  |  | Gbp CDS R | cgagcccttgctagcCTGGTCGATTCTAACCAATATC |  |  |
|  |  | pLinker-Tyx3 F | GTTAGAATCGACCAGgctagcaagggctcgggc | Amplify pLinker-Tyx3 region from GP-mAID-3HA-TIR1-Nluc-P2A-neo plasmid for gibson assembly | [4]; IDT |
|  |  | pLinker-Tyx3 R | TTGGTAAAAAAAACATTAGTCGAGCGGGTCCTGG |  |  |
|  |  | CenH3 3'UTR F | GGACCCGCTCGACTAATGTTTTTTTTACCAAATAAATAGCTCCTTATTTGC | Amplify CENH3 3'UTR region from TK-3HA-CENH3-Nluc-P2A-neo-TK plasmid (not featured in MS) for gibson assembly |  |
|  |  | CenH3 3'UTR R | ttagtttccccacctCCGCTACTACCGACCGTTTATTG |  |  |
|  |  | P2A Linker | AGACTTTATTggaagcggagctactaacttcagcctgctgaagcaggctggagacgtggaggagaaccctggacctATGGTATCTG | P2A Linker gBlock for for gibson assembly |  |
|  | Gene insertion confirmation | Cp-TK F2 | ACCATCTAACACAAAGAATTCAACA | To check correct gene insertion from DNA purified from mouse feces. | [2]; IDT |
|  |  | 3HA R | ggcataatctggaacatcgtaagg |  |  |
|  |  | Neo F1 | GCTGAAGAACTTGGTGGTGA |  |  |
|  |  | Cp TK-R | CCTACCATAAGCATCACCAGT |  |  |
| Epitope tagging of cgd4_2030 (CENH3) (spaghetti monster HA) | CENH3 (cgd4_2030)-smHA-Nluc-P2A-neo | Linker-F1 | gctagcaagggctcgggc | Amplify spaghetti monster tag region from p2870-smHA-Nluc-P2A-neo | [2]; [4]; IDT |
|  |  | AldoUTR_R | ggcgcgccaaataaagtaaagtttatcg |  |  |
|  |  | Aldo 3UTR-F | atgcatcttcatttagtatcttaggt | Amplify CENH3 (cgd4_2030)-3HA-Nluc-P2A-neo without the 3HA tag |  |
|  |  | pLinker-R2 | cagctgggtcgagcccga |  |  |
|  | Gene insertion confirmation | CenH3 5'UTR check F2 | AGTAGCCCCATTAGCCCAAG | To check correct gene insertion from DNA purified from mouse feces. | [2]; IDT |
|  |  | pLinker-R2 | cagctgggtcgagcccga |  |  |
|  |  | CenH3 3'UTR check R | GGCGGGGATAATTCATGTTT |  |  |
|  |  | NeoF1 | GCTGAAGAACTTGGTGGTGA |  |  |
